# Cytosolic heme metabolism by alternative localization of heme oxygenase 1 in plant cells

**DOI:** 10.1101/2023.10.30.564862

**Authors:** Yingxi Chen, Kohji Nishimura, Mutsutomo Tokizawa, Yoshiharu Y. Yamamoto, Yoshito Oka, Tomonao Matsushita, Kousuke Hanada, Kazumasa Shirai, Shoji Mano, Takayuki Shimizu, Tatsuru Masuda

## Abstract

Heme, an organometallic molecule, is widely engaged in oxygen transport, electron delivery, enzymatic reactions, and signal transduction. Additionally, heme serves as a precursor to phytochromobilin, the chromophore of plant phytochrome. Heme oxygenase (HO) initiates the first committed step in heme metabolism. Our transcription start site-sequencing (TSS-seq) revealed that *HO1* in *Arabidopsis thaliana* and *Oryza sativa* (rice) has two TSSs, producing long (*HO1L*) and short (*HO1S*) transcripts, with or without an intact N-terminal plastid transit peptide. *HO1L* and *HO1S* products localize in plastids and the cytosol, respectively. In Arabidopsis, *HO1L* is prevalent in light-exposed shoots, while *HO1S* is clearly detected in roots and etiolated seedlings. During de-etiolation and early development, *HO1L* ratio gradually rises and *HO1S* ratio decreases. Light perception via phytochrome and cryptochrome elevates *HO1L* ratio and reduce *HO1S* ratio through the functioning of HY5 and HYH transcription factors, and the suppression of DET1, E3 ubiquitin ligase COP1, and PIFs transcription factors. As expected, *HO1L* product was able to complement the *HO1*-deficient mutant *gun2* (*hy1*), but surprisingly, *HO1S* expression could also restore the short hypocotyl phenotype and high pigment content, and make the mutant recover from the *gun* phenotype. This indicates the formation of functional holo-phytochrome within these lines. Our work highlights the presence of a cytosolic pathway for heme metabolism, especially during etiolation and early development. Furthermore, it supports the hypothesis that a mobile heme signal is involved in the mediation of retrograde signaling from the chloroplast.

**Significance Statement:** In this research, through both TSS-seq and CAGE-seq, we discovered that the *HO1* (*GUN2* or *HY1*) gene in both Arabidopsis and rice has two TSSs, generating *HO1L* and *HO1S* transcripts. We reveal that the *HO1* TSS regulation pathway is the same as the light signaling pathway. Significantly, our study identifies that a cytosolic heme metabolism pathway is existent in plant cells.

## Introduction

In plants, chloroplast biogenesis encompasses a series of stages that commence with the development of proplastids (1). To develop into chloroplasts, proplastids in the photosynthetic mesophyll cells of a leaf import various proteins from the cytosol. This process involves protein targeting sequences that direct proteins to the correct location within the developing organelle. In dark-grown plants, the mesophyll cell proplastids develop into etioplasts. When etiolated plants are transferred to the light, etioplasts differentiate into chloroplasts. Thus, both environmental signals, such as light and intrinsic developmental signals, determine the fate of the undifferentiated proplastids.

For sensing light signals, phytochrome and cryptochrome are well-known photoreceptors in plants that regulate various aspects of plant growth and development in response to light conditions (2, 3). Phytochrome primarily responds to red and far-red lights, whereas cryptochrome predominantly reacts to the blue light. Phytochrome is composed of an apoprotein to which a linear tetrapyrrole chromophore, phytochromobilin (PΦB), is covalently attached. It exists in either of two photoconvertible forms: the red light-absorbing (P_r_) or the far-red light-absorbing (P_fr_) form. Recent research reveals that light signaling via phytochrome prompts the selection of alternative promoters, enabling plants to adapt their protein localizations in response to changing light conditions (4).

Heme is a crucial molecule found in various organisms, including animals, plants, and microorganisms. It is a ferrous iron (Fe^2+^) complex coordinated within a porphyrin ring structure and functions as a prosthetic group incorporated into various apoproteins. Heme serves multiple important functions in various biological processes, including binding and transport of oxygen, electron transfer, enzymatic reactions, signal transduction, and gene expression. In plants, heme is crucial for not only photosynthesis, but also photomorphogenesis, serving as a precursor of PΦB. In particular, heme is also proposed to be a biogenic retrograde signal which transduces information from plastid to the nucleus (5). Despite the importance of heme, its accumulation can be toxic to cells due to its pro-oxidant and pro-inflammatory properties (6). Thus, organisms, including plants, have evolved mechanisms to regulate heme levels for a balanced supply of various cellular functions.

Heme is synthesized through a conserved biosynthetic pathway which starts from a precursor molecule 5-aminolevulinic acid (ALA) (7). ALA is then processed through a series of enzymatic steps, including the addition of extra pyrrole rings and decarboxylation to form protoporphyrin IX (Proto-IX) (8). Once Proto-IX is synthesized, it serves as a critical precursor for both heme and chlorophyll (Chl) syntheses. In the heme branch, the ferrochelatase (FC) inserts Fe^2+^ into the Proto-IX ring. In general, plants have two genes encoding FC (*FC1* and *FC2*), which show differential tissue-specific and development-dependent expression profiles, such that *FC2* is light-dependent and mainly expressed in photosynthetic tissues, whereas *FC1* is stress-responsive and ubiquitously expressed in all tissues (9). In mammals and yeast, heme is synthesized in the mitochondrion, whereas in plant cells, it is biosynthesized and metabolized only in the plastid (10). Moreover, heme is transported to the cytosol and various other organelles, including mitochondrion, peroxisome, vacuole, and endoplasmic reticulum (11).

Heme oxygenase (HO) is the key enzyme involved in heme metabolism, and it is highly conserved across mammalian, plant, and fungal kingdoms (12, 13). The HO enzyme cleaves the heme molecule to release biliverdin IXα (BV-IXα), carbon monoxide, and Fe^2+^ (14, 15). In plants, this enzymatic reaction is essential for maintaining cellular homeostasis and regulating various physiological processes such as chloroplast biogenesis, photomorphogenesis, antioxidant defense, iron mobilization, and jasmonic acid signaling (12, 16). Significantly, BV-IXα, which serves as the precursor to PΦB, plays a pivotal role in the perception of light (17). Arabidopsis HO family contains four members: HO1 (also known as GENOMES UNCOUPLED 2, GUN2, or HY1), HO2, HO3, and HO4 (18–20). HO1, HO3, and HO4 serve in the first committed step for heme metabolism, and among them, HO1 plays the fundamental role. All HO isoforms are transported to plastids via their N-terminal transit peptides (TPs). BV-IXα is further converted to PΦB by PΦB synthase HY2 (also known as GUN3) (21). It is reported that HO1 and HY2 are ferredoxin-dependent reductases, obtaining electrons from photosynthetic electron transport in chloroplast (22, 23). Furthermore, since HY2 is also considered to be localized in plastids, it has been believed that PΦB synthesis takes place only in plastids (23).

Here, through transcription start site-sequencing (TSS-seq), we identified two TSSs of the *HO1* gene, which generated a long mRNA (designated *HO1L*) containing the intact plastid TP and a short one (designated *HO1S*) lacking the TP. Our study identifies that the *HO1* TSSs regulation pathway is the same as the light signaling pathway. In detail, the upregulation of *HO1L* ratio and the downregulation of *HO1S* ratio are in accordance with the photomorphogenesis signaling pathway. Moreover, in this study, we showed that *HO1L* and *HO1S* products were localized in plastids and the cytosol, respectively, through subcellular localization analysis. Complementation assay demonstrated that not only *HO1L* but also *HO1S* product restored from phenotypes of the *HO1* deficient mutant *gun2-1*. Especially, besides the *HO1L* transgenic line, the *HO1S* transgenic line discards the *gun* phenotype, consistent with the hypothesis that FC1-derived heme functions as a positive plastid signal for the expression of *photosynthesis-associated nuclear genes* (*PhANGs*). Our study defies the common sense that PΦB biosynthesis takes place only in plastids. The identification of a cytosolic heme metabolism/PΦB biosynthesis pathway has novel implications for understanding the heme metabolism and its regulation in plants.

## Results

### *HO1* has two TSSs in both Arabidopsis and rice

In this study, we employed the TSS-seq screening to investigate genes exhibiting multiple TSSs in both Arabidopsis and rice. We compared genes with different TSSs in light-grown shoots and roots, and etiolated seedlings of Arabidopsis, and light-grown shoots and roots of rice. As a result, we found overlapping 19 gene families containing 24 genes of Arabidopsis and 27 genes of rice to have different TSSs. Among them, we identified *HO1* as a gene that alters TSSs in response to light in both Arabidopsis and rice, suggesting that the *HO1* gene is regulated by promoter switching in plants. Taking Arabidopsis as an example, in shoots of seedlings exposed to the white light, the *HO1* gene is mainly transcribed into *HO1L*, the long mRNA, containing the intact plastid TP, whereas in roots and etiolated seedlings, it is also transcribed into *HO1S*, the short one, with the truncated TP (Fig. 1a, b). Furthermore, cap analysis of gene expression-sequencing (CAGE-seq) confirmed that *HO1* gene in both Arabidopsis and rice has different TSSs (*SI Appendix,* Fig. S1a, b). Therefore, it is clarified that *HO1* has two TSSs which are regulated in both light- and organ-dependent manner. We also examined whether other *HO* genes and genes involved in the heme branch (*FC1, FC2,* and *HY2*) have multiple TSSs or not (*SI Appendix,* Fig. S1c to h). However, no gene has multiple TSSs which shows that the TSS regulation is specific to the *HO1* gene in the heme branch.

**Figure 1.**
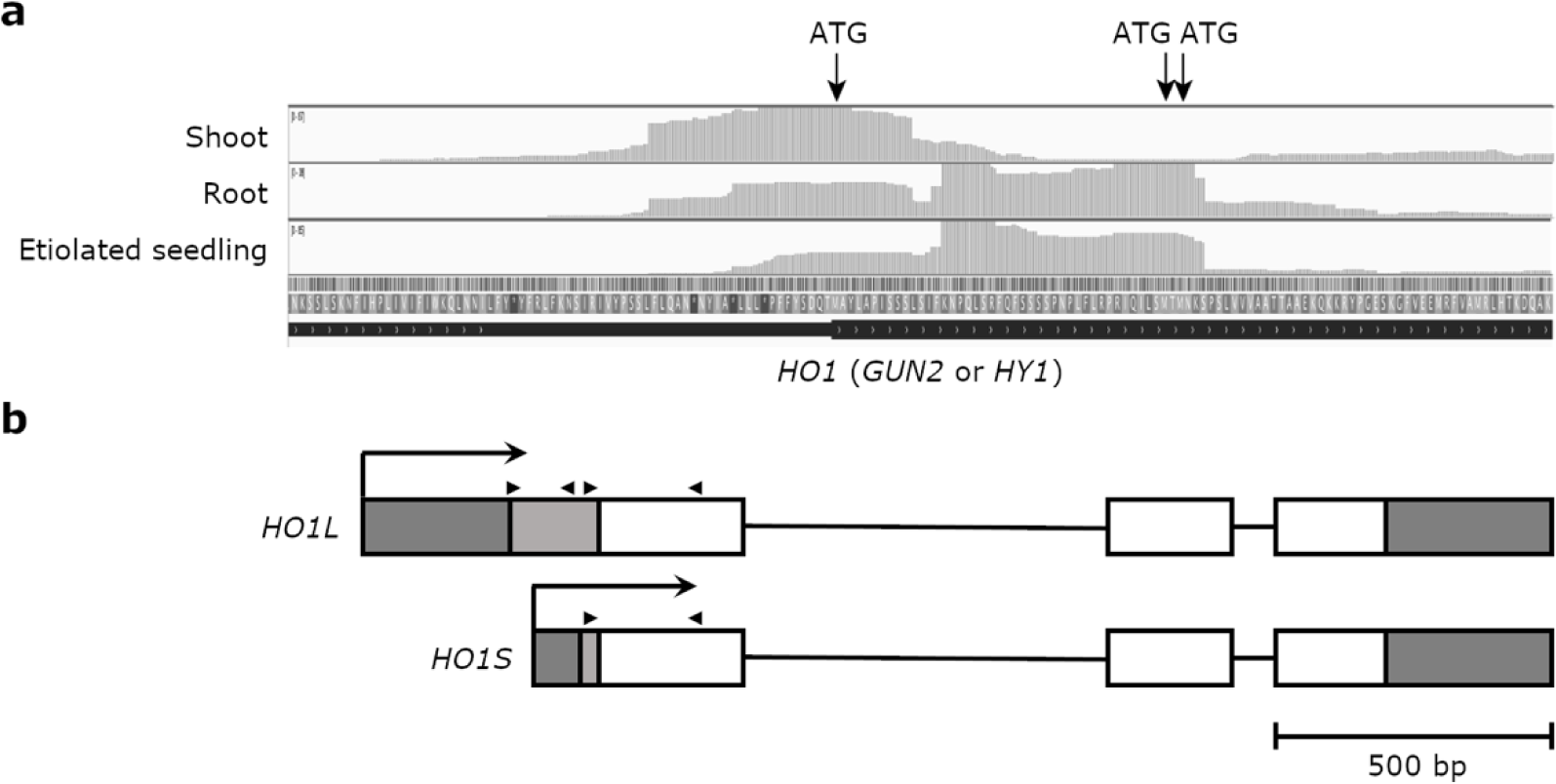
TSS-seq and TSSs of the *HO1* gene. a, Screenshot of the representative Arabidopsis *HO1* TSS-seq data. The upper panels are TSS signals of shoot, root, and etiolated seedling, respectively. The middle and bottom panels are the N-terminal amino acid sequence of HO1 and the N-terminal of the *HO1* gene structure. ATG refers to the translation start codon. b, Schematic representation of *HO1* pre-mRNAs in Arabidopsis cells. The diagram is constructed based on the data from Phytozome 13 (https://phytozome-next.jgi.doe.gov), UniProt (https://www.uniprot.org/), and sequencing results of 5’ RACE-PCR. Dark gray box and black line illustrate untranslated region (UTR) and intron, respectively. Both light gray and white boxes represent exons, with the light gray box specifically indicating the plastid TP. Black arrow illustrates the presumed TSS. Black arrowhead illustrates positions of primers utilized for the 5’ RACE-qPCR. Scale bar, 500 bp.

To check whether *HO1L* product containing the intact plastid TP is transported into plastids and *HO1S* product lacking the TP retains in the cytosol, we performed subcellular localization analysis. To avoid the promoter switching of own *HO1* promoter, *cauliflower mosaic virus 35S promoter* (*p35S*) was utilized. Vectors harboring genes encoding Arabidopsis *HO1L* or *HO1S* fused to *GFP*, *mClover3* (24) under the control of *p35S*, *p35S*::*HO1L*-*mClover3* and *p35S*::*HO1S*-*mClover3*, were introduced into *Pisum sativum* (pea) and *Nicotiana benthamiana* (tobacco) leaves by particle bombardment. In *HO1L*-introduced leaves, we detected clear fluorescent signals that overlaying those of a chloroplast marker, plastid TP of a small subunit of Arabidopsis RIBULOSE BISPHOSPHATE CARBOXYLASE SMALL CHAIN 1A (AtRBCS1A-TP) (25) fused RFP, mScarlet (26) signal, and Chl autofluorescence in pea and tobacco (*SI Appendix,* Fig. S2). While dispersed signals overlaying the cytosol-derived mScarlet fluorescence were detected in *HO1S*-introduced leaves. These results show the plastidial localization of the *HO1L* product and cytosolic localization of the *HO1S* product.

### *HO1* TSSs are regulated by light and development

To explore how light regulates *HO1* TSSs, we examined de-etiolated seedlings of wide-type Arabidopsis Columbia-0 (Col-0) which was first grown under the dark for four days and then transferred to the white light for different hours (Fig. 2a). mRNA was extracted from the whole seedling of all the samples, reverse-transcribed with the 5’ Rapid Amplification of cDNA Ends (RACE), and analyzed by PCR with assigned primers (*SI Appendix,* Table S1). Results show that ratios of *HO1L* and *HO1S* are stable within the first six hours after being transferred to the white light; yet with the extension of light exposure time, the *HO1L* ratio increases and the *HO1S* ratio decreases. To determine *HO1L* and *HO1S* ratios more precisely, we developed the 5’ RACE-qPCR method using two sets of primers to measure transcription levels of *HO1L* and total *HO1* (*SI Appendix,* Table S1). After determination of each transcript level, *HO1S* level was calculated by subtracting *HO1L* from total *HO1* level. In the early developmental stage of young seedlings grown under continuous white light, the *HO1L* ratio gradually increases and the *HO1S* ratio decreases (Fig. 2b), suggesting that *HO1S* expression prevailed in the early stage of development. These data uncover that the *HO1* TSS slowly responds to white light signals.

**Figure 2.**
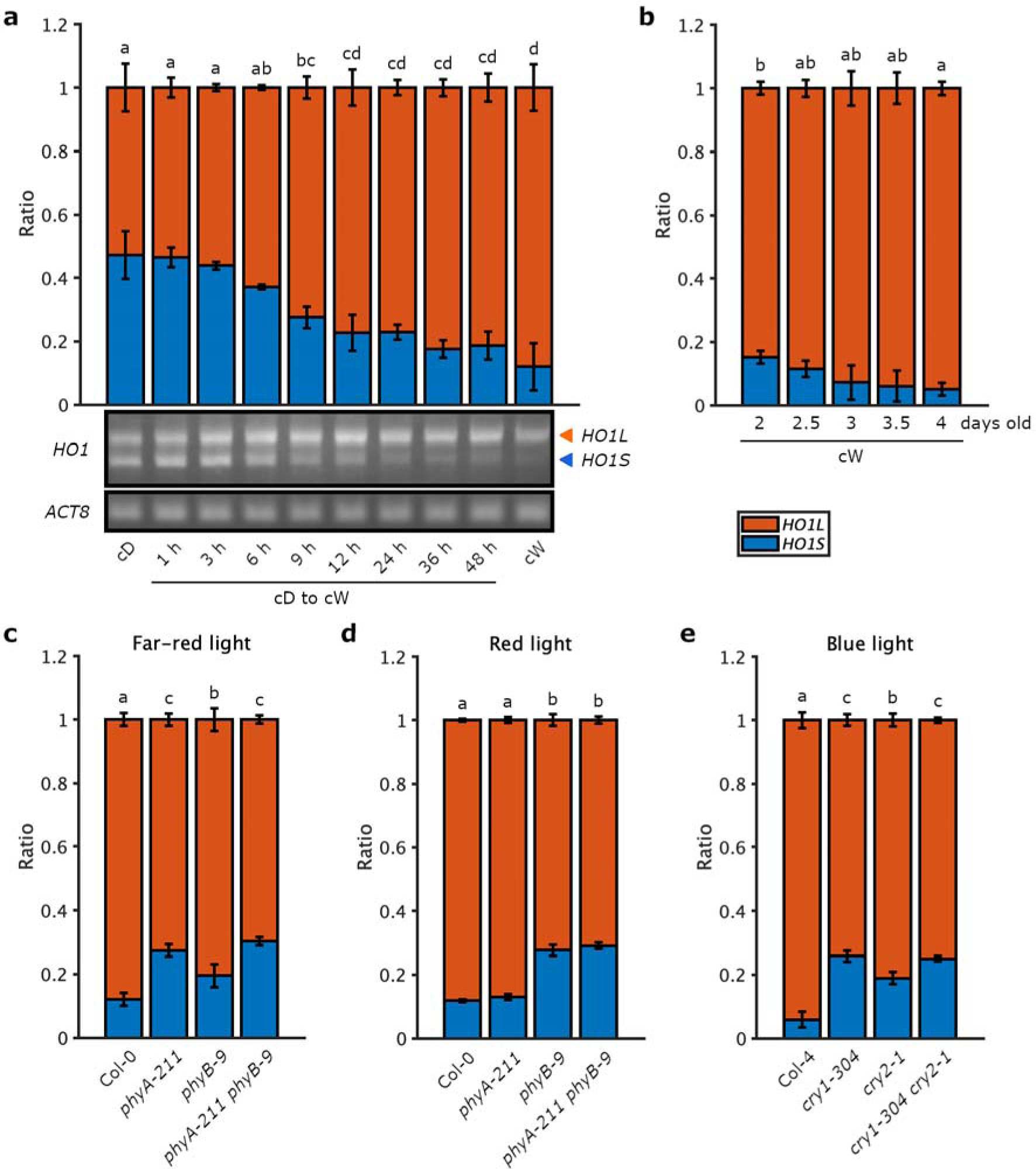
*HO1L* and *HO1S* ratios in seedlings cultivated under various light conditions. a, *HO1L* and *HO1S* ratios in de-etiolated seedlings. Wide-type Col-0 was cultivated under the dark for four days before being irradiated with the white light (55 μmol photons m^-2^ s^-1^) for indicated periods. Four-day-old seedlings cultivated under the continuous white light (cW) or dark (cD) were used as controls. The gene specific primer (GSP) of *HO1*, GSP-*HO1*, was used in the 5’ RACE-PCR. PCR products were loaded onto a 2.5% DNA gel for electrophoresis. Lower are representative pictures of the gel electrophoresis. Orange and blue arrowheads illustrate *HO1L* and *HO1S* bands, respectively. The upper bar graph was obtained by analyzing the lower *HO1* gel picture with the CSAnalyzer4 software. *ACT8*, *Actin 8*, is the internal control, which is amplified with *ACT8*-F and *ACT8*-R. Primers used in PCR are listed in *SI Appendix*, Table S1. Two biological replicates were performed, and the representative one is shown. The data mean ± SD was calculated from values of three technical replicates. MATLAB software was used to analyze the data. p<0.05, one-way ANOVA, and Turkey’s multiple comparison tests. b, *HO1L* and *HO1S* ratios in early developmental seedlings. Col-0 was cultivated under the cW for indicated periods. Two pairs of primers, *HO1L*-F and *HO1L*-R, and *HO1*-F and *HO1*-R were used in the 5’ RACE-qPCR for measuring Ct values of *HO1L* and total *HO1*. Primers used in qPCR are listed in *SI Appendix*, Table S1. At least three biological replicates were performed, and the representative one is shown. c to e, 5’ RACE-qPCR analysis of *HO1L* and *HO1S* ratios in seedlings grown under different light colors. Col-0, *phyA-211*, *phyB-9*, and *phyA-211 phyB-9* were cultivated under the continuous far-red (5 μmol photons m^-2^ s^-1^) (c) or red (15 μmol photons m^-2^ s^-1^) (d) light for four days. Col-4, *cry1-304*, *cry2-1*, and *cry1-304 cry2-1* were cultivated under the continuous blue (15 μmol photons m^-2^ s^-1^) light (e).

There are two major molecular species of phytochrome, phytochrome A (PHYA) and phytochrome B (PHYB) (27), and two cryptochromes, cryptochrome 1 (CRY1) and cryptochrome 2 (CRY2) in Arabidopsis. To investigate the specific light signal and photoreceptor involved in the mediation of *HO1* TSS, we examined *HO1L* and *HO1S* ratios in Arabidopsis phytochrome or cryptochrome deficient mutants grown under continuous monochromatic illumination of far-red, red, or blue light wavelengths (Fig. 2c to 2e). Under the far-red light, *phyA-211* and *phyA-211 phyB-9* accumulated less *HO1L* and higher *HO1S* ratios than Col-0. Under the red light, *phyB-9* and *phyA-211 phyB-9* accumulated less *HO1L* and higher *HO1S* ratios than Col-0. Under the blue light, Arabidopsis wide-type Columbia-4 (Col-4) based *cry1-304*, *cry2-1*, and *cry1-304 cry2-1* accumulated less *HO1L* and higher *HO1S* ratios than the wide-type. Taken together, these lines of evidence clearly clarify that both phytochrome and cryptochrome absorbing far-red, red, and blue lights regulate the *HO1* TSS.

### Central light repressors and transcription factors related to light signaling are involved in *HO1* TSSs regulation

CONSTITUTIVE PHOTOMORPHOGENESIS 1 (COP1) prevents the expression of light-responsive genes by ligating specific transcription factors such as ELONGATED HYPOCOTYL 5 (HY5) with ubiquitin molecules, marking them for degradation by the 26S proteasome (28). HY5 and HY5 HOMOLOG (HYH) promote photomorphogenesis by activating the expression of genes involved in Chl biosynthesis, light harvesting, and other light-responsive processes (29). They also regulate the expression of genes involved in skotomorphogenesis. DE-ETIOLATED 1 (DET1) forms a complex with other proteins, including the COP1 (30), and acts downstream of photoreceptors, such as phytochrome and cryptochrome, in light signaling pathways. 5’ RACE-qPCR discovered that *cop1-4* and *det1-1* accumulated significantly higher *HO1L* and lower *HO1S* ratios than Col-0 under the dark, indicating that both COP1 and DET1 play significant roles in the *HO1* TSS regulation (Fig. 3a, b). 5’ RACE-qPCR results showed that under the white light, *hy5-215* and *hy5-2 hyh* accumulated less *HO1L* and more *HO1S* ratios than Col-0 (Fig. 3c and *SI Appendix,* Fig. S3a).

**Figure 3.**
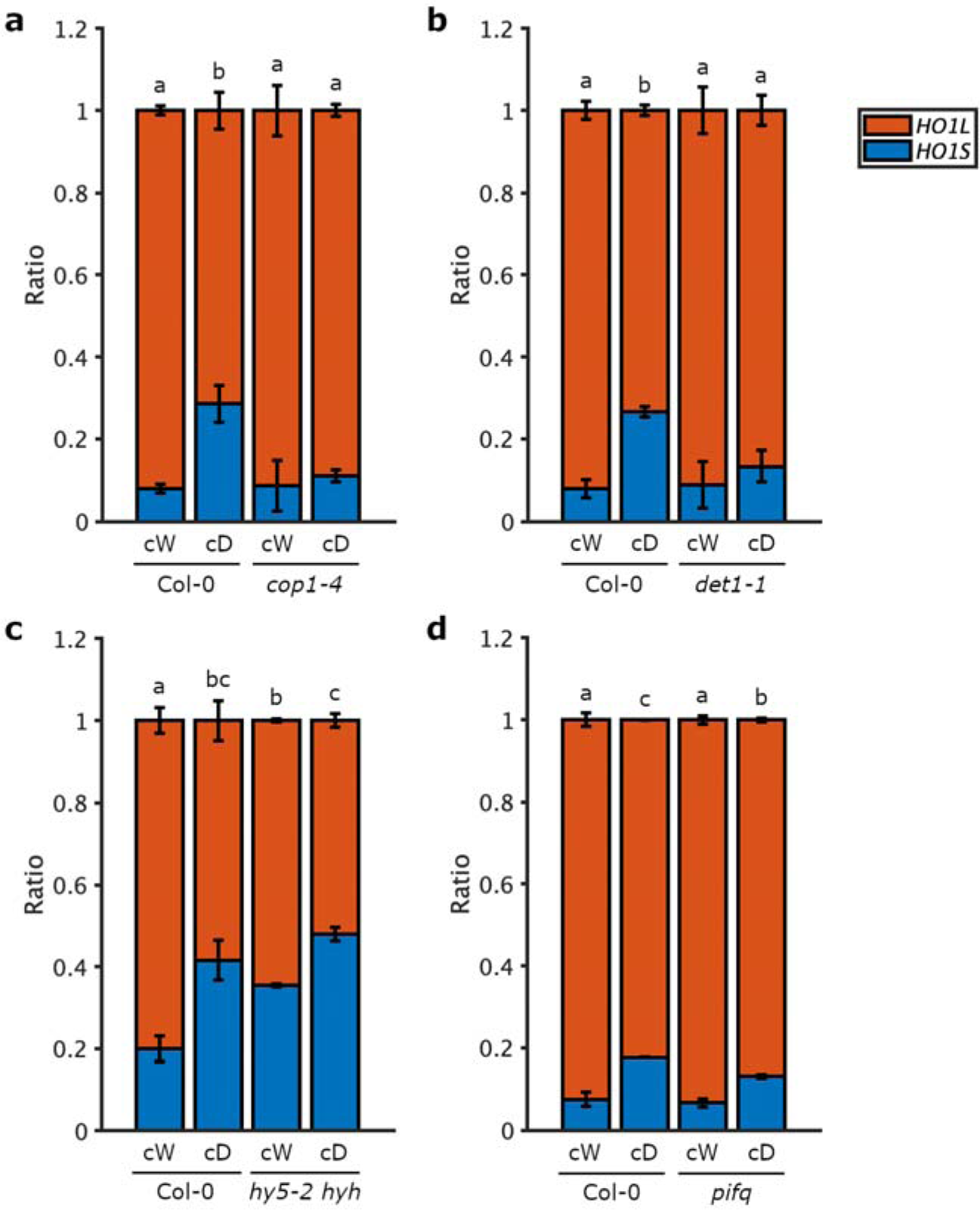
5’ RACE-qPCR analysis of *HO1L* and *HO1S* ratios in mutants of light regulated factors. a, *HO1L* and *HO1S* ratios in the COP1 mutant. Col-0 and *cop1-4* were cultivated under the continuous white light (cW, 55 μmol photons m^-2^ s^-1^) or dark (cD) for four days. b, *HO1L* and *HO1S* ratios in the DET1 mutant. Col-0 and *det1-1* were cultivated under the cW or cD condition for four days. c and d, *HO1L* and *HO1S* ratios in mutants of transcription factors. Col-0, *hy5-2 hyh* (c), and *pifq* (d) were cultivated under the cW or cD condition for four days.

The PHYTOCHROME INTERACTING FACTOR (PIF) family in Arabidopsis consists of eight members (31). They act as central regulators that integrate and transmit light signals to downstream gene expression networks. We found that the PIF quadruple mutant *pifq* (*pif1*/*pif3*/*pif4*/*pif5*) accumulated more *HO1L* and less *HO1S* ratios under the dark condition than Col-0 (Fig. 3d). These data indicate that key transcription factors of HY5 and HYH, which are regulated in the opposite way by light repressors COP1 and DET1, and transcription factors PIFs are involved in the *HO1* TSS regulation. Although it seems that the effect of PIFs on the *HO1* TSS is much less than those of COP1, DET1 and HY5/HYH.

### *HO1* TSS regulation is not related to the chloroplast function

In non-photosynthetic organ roots, higher level of *HO1S* transcript was detected (Fig. 1a). In addition, the light-dependent TSS regulation of the *HO1* gene was slower than other light-responsive gene transcription responses, which becomes apparent after 6 h of illumination during de-etiolation (Fig. 2a). Therefore, we hypothesized that the *HO1* TSS regulation may be related to the chloroplast development rather than the direct promoter switching by transcription factors. It is reported that herbicides such as norflurazon (NF), an inhibitor of the carotenoid biosynthesis enzyme phytoene desaturase, or the plastid translation inhibitor, lincomycin (Lin), disrupt the chloroplast function (32, 33). We treated Col-0 with NF or Lin and analyzed in light-grown shoots and roots, and etiolated seedlings. However, these treatments did not affect the *HO1L* and *HO1S* ratios in all samples (*SI Appendix,* Fig. S3b, c), showing that chloroplast functionality is not related to the TSS regulation.

### Complementation analysis of *HO1L* and *HO1S* transgenic lines in *gun2-1*

The *ho1* mutant (*gun2-1* or *hy1-1*) is reported to show reduced light responses, long hypocotyl, incomplete chloroplast development with reduced Chl accumulation, and less extensive grana thylakoid membrane stacking (34). Here, we showed that *HO1* transcripts of both *HO1L* and *HO1S* are present in Arabidopsis and rice, and the *HO1L* product is localized in plastids (*SI Appendix,* Fig. S2). However, the function of the cytosol localized *HO1S* product is unclarified. To explore functions of the *HO1S* product, we constructed *HO1L* and *HO1S* transgenic lines in the *gun2-1* background. The same vectors utilized in the upward transient expression assay were transformed into Col-0 based *HO1* deficient mutant *gun2-1* by agrobacterium-mediated floral dip transformation, generating *p35S*::*HO1L*-*mClover3*/*gun2-1* and *p35S*::*HO1S*-*mClover3*/*gun2-1* (hereafter referred to as *HO1L*/*gun2-1* and *HO1S*/*gun2-1*) lines. Expression levels of *HO1L* and *HO1S* were measured by both RT-qPCR and western blot (Fig. 4a, b). *HO1* transcription levels in the 7-1 *HO1L*/*gun2-1* and 4-5-3 *HO1S*/*gun2-1* strains are 62.5 and 11.0 times higher than that in the wild type Col-0, respectively.

**Figure 4.**
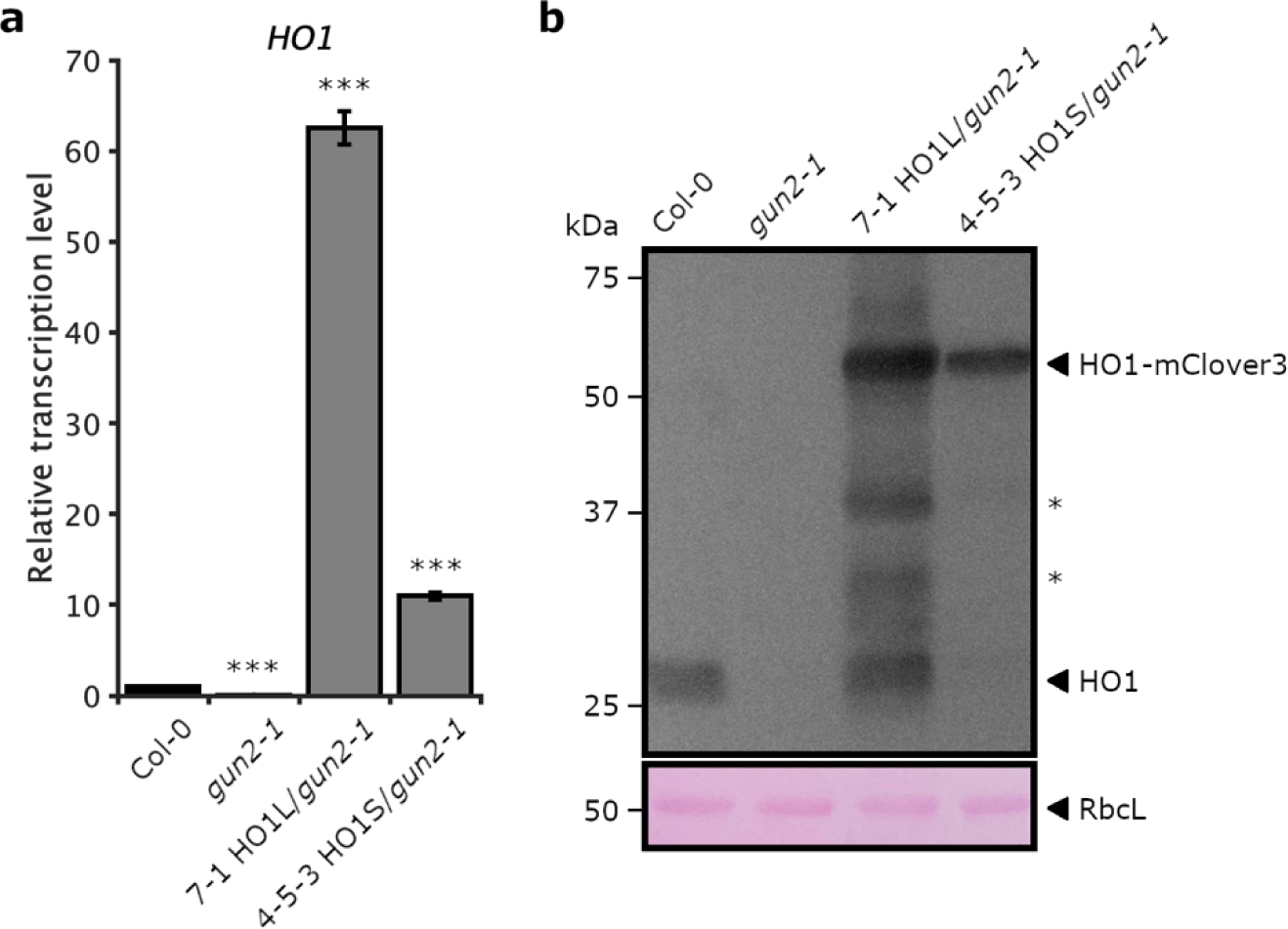
The construction of HO1L and HO1S transgenic lines. a, RT-qPCR analysis of *HO1* transcription levels. Seedlings were grown under the continuous white light (cW, 55 μmol photons m^-2^ s^-1^) for four days. *HO1*-F and *HO1*-R were used for measuring transcription levels of *HO1* including both *HO1L* and *HO1S*. *ACT8* was the internal control. Primers used in qPCR are listed in *SI Appendix*, Table S1. At least three biological replicates were performed, and the representative one is shown. *** indicates significant differences at p<0.005 according to student’s t-test. b, Western blot analysis of HO1 expression levels. Seedlings were cultivated under the cW condition for four days. * indicates non-specific bands. Same PVDF membrane was stained by Ponceau S to display protein RbcL, a large subunit of RuBisCO, as the loading control.

Consistent with the previous study (34), *gun2-1* showed yellow cotyledons in four-day-old strains cultivated under continuous white light (Fig. 5a). It was intriguing that like the 7-1 *HO1L*/*gun2-1* strain, the 4-5-3 *HO1S*/*gun2-1* line showed green cotyledons. To check phenotypes of the *HO1S* transgenic line, the vector harboring *p35S*::*HO1S*-*mClover3* was also transformed into Arabidopsis Landsberg erecta (Ler) based *HO1* deficient mutant *hy1-1*, generating *p35S*::*HO1S*-*mClover3*/*hy1-1* (hereafter referred to as *HO1S*/*hy1-1*) transgenic lines. Consistent with the phenotype of 4-5-3 *HO1S*/*gun2-1* seedling, 6-1, 8-3, and 8-5 *HO1S*/*hy1-1* transgenic lines showed green cotyledons (*SI Appendix,* Fig. S4a). These data indicates that *HO1S* product has significant complementary function for the *HO1* deficiency in respect to plant growth and development.

**Figure 5.**
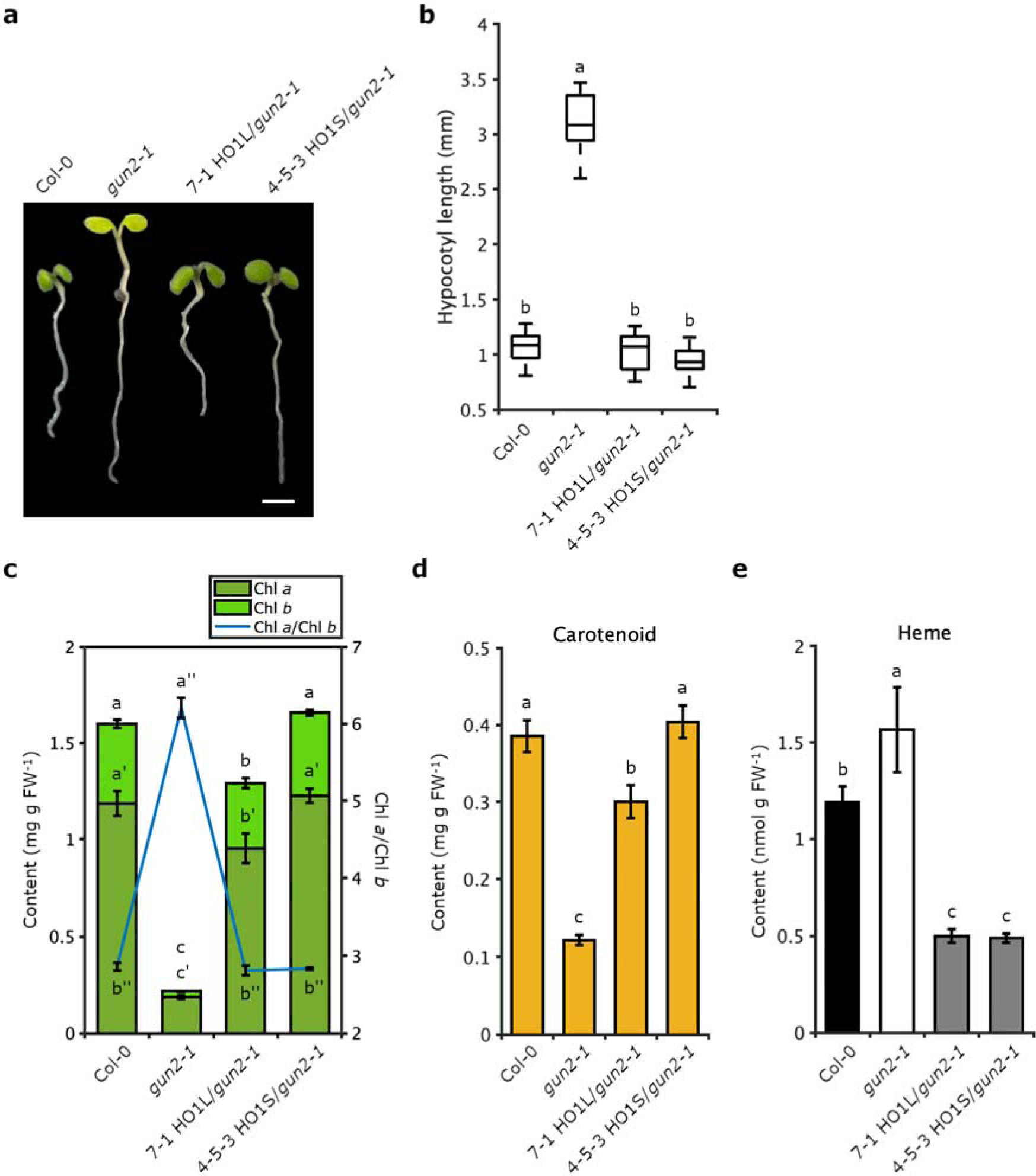
Hypocotyl length and pigment contents of HO1L and HO1S transgenic lines. a, Phenotype of four-day-old transgenic lines. Seedlings were cultivated under the continuous white light (cW, 55 μmol photons m^-2^ s^-1^) for four days. Scale bar, 2 mm. b, Hypocotyl length of transgenic lines. Strains were cultivated under the cW condition for four days. Hypocotyl lengths were measured using ImageJ, and measurements were taken with 20 seedlings from each line. p<0.05, one-way ANOVA, and Turkey’s multiple comparison tests. c to e, Chl content (c), Chl *a*/Chl *b* ratio (c), carotenoid content (d), and free heme level (e) of transgenic lines. Seedlings were grown under the cW condition for four days. The data mean ± SD was calculated from values of three technical replicates.

### Explore the HO1S function

To confirm the localization of *HO1L* and *HO1S* products, the fluorescence microscopic imaging was performed using protoplasts prepared from well-expanded leaves of *HO1L*/*gun2-1* and *HO1S*/*gun2-1* lines. Consistent with the transient expression assay (*SI Appendix,* Fig. S2), *HO1L*-*mClover*3 and *HO1S-mClover3* products were localized in the chloroplast and cytosol, respectively, in the transgenic lines (Fig. 6).

**Figure 6.**
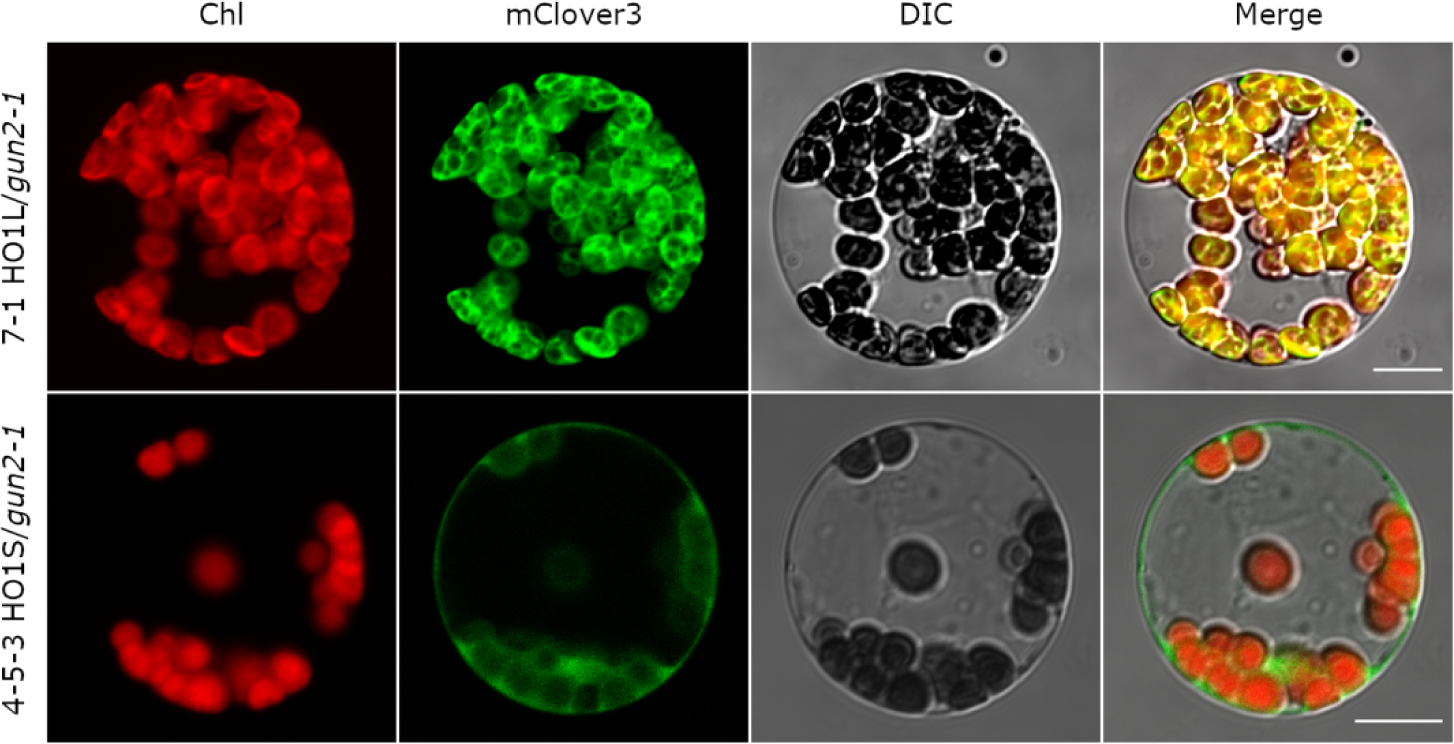
Localizations of HO1L-mClover3 and HO1S-mClover3 in transgenic lines. Leaves of three-week-old strains grown under the short-day condition (SD, 8 h 120 μmol photons m^-2^ s^-1^ white light/16 h dark) were detached for the protoplast isolation. At least three biological replicates were performed, and around 10 protoplasts of each biological replicate were observed under the confocal laser-scanning microscope (CLSM). Single-section images of the Chl channel (red), the GFP channel (green), DIC (gray), and their merge in each panel are shown. Bars indicate 10 µm.

In *gun2-1*, because of the lack of phytochrome chromophore PΦB, phytochrome is dysfunctional and the photomorphogenesis process is affected, resulting in elongated hypocotyl length (Fig. 5a, b). In the complementation lines, not only 7-1 *HO1L*/*gun2-1* but also 4-5-3 *HO1S*/*gun2-1* recovered the short hypocotyl length as observed in Col-0. Similarly, 6-1, 8-3, and 8-5 *HO1S*/*hy1-1* transgenic lines recovered the short hypocotyl length as observed in Ler (*SI Appendix,* Fig. S4a, b). Contents of pigments including Chl *a*, Chl *b*, and carotenoid were also restored the low pigment levels in *gun2-1*. Contents of these pigments in 7-1 *HO1L*/*gun2-1* and 4-5-3 *HO1S*/*gun2-1* were a slightly lower and the same as those in Col-0, respectively (Fig. 5c, d). Similarly, 6-1, 8-3, and 8-5 HO1S/*hy1-1* lines showed slightly lower levels than Ler (*SI Appendix,* Fig. S4c, d). Furthermore, the Chl *a*/Chl *b* ratio in all complemented lines of *HO1L*/*gun2-1*, *HO1S*/*gun2-1*, and *HO1S*/*hy1-1* returned to the wild-type level (Fig. 5c, *SI Appendix,* Fig. S4c). It is of note that total free heme content in *gun2-1* was higher than that in wild-type, while those in 7-1 *HO1L*/*gun2-1* and 4-5-3 *HO1S*/*gun2-1* lines were much lower than that in Col-0 (Fig. 5e), suggesting enhanced heme degradation in plastids and the cytosol, respectively, in transgenic lines.

Western blot analysis of PHYA and PHYB showed that both proteins were highly accumulated in the *gun2-1* mutant under the light condition (Fig. 7a), consistent with the fact that *gun2-1* contains dysfunctional apo-phytochromes (35). Whereas PHYA and PHYB amounts in both 7-1 *HO1L*/*gun2-1* and 4-5-3 *HO1S*/*gun2-1* transgenic lines were decreased to the wild-type level, suggesting that functional holo-phytochrome is synthesized in these transgenic strains.

**Figure 7.**
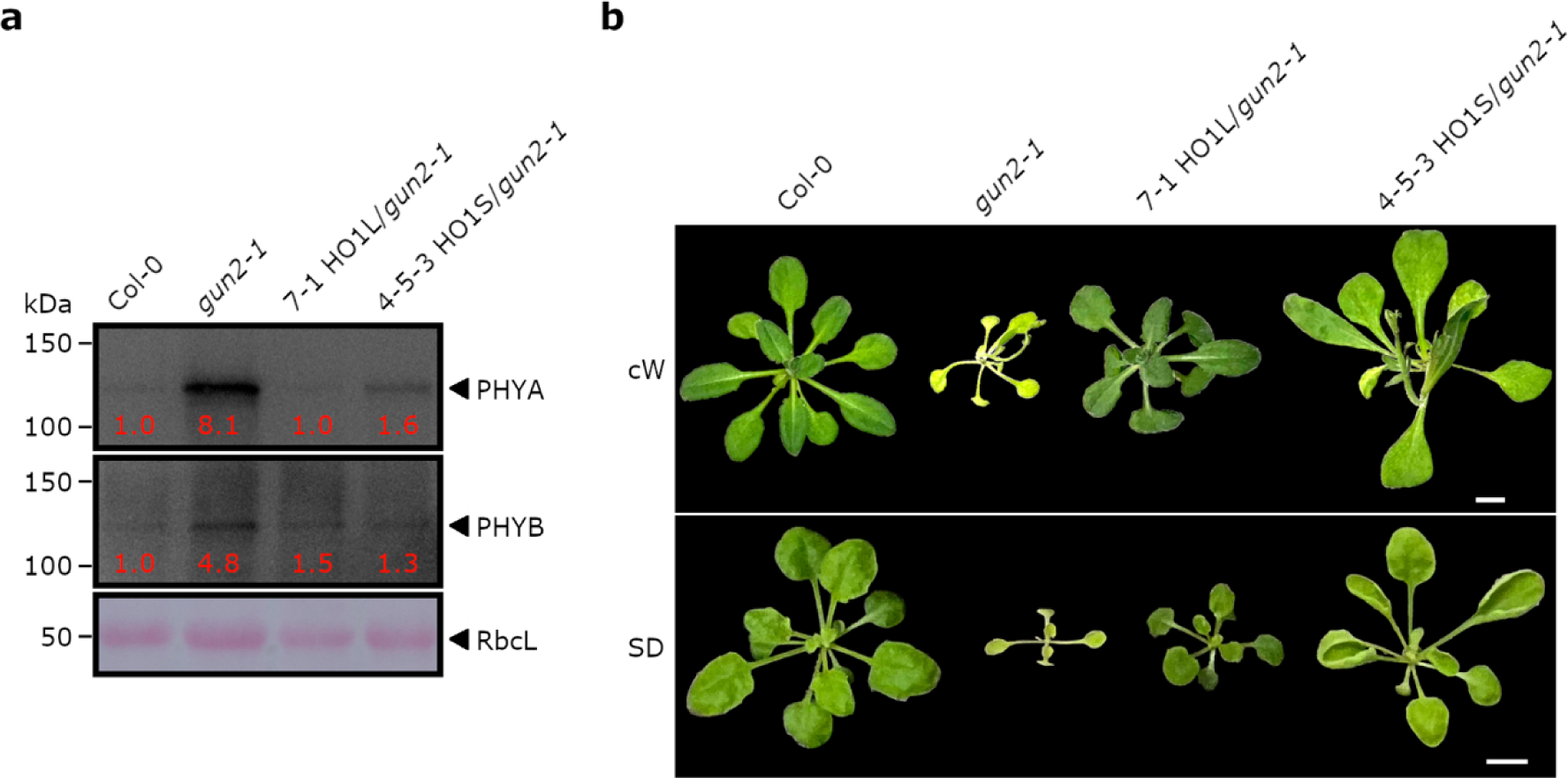
Phytochrome expression and phenotype of HO1L and HO1S adults. a, Western blot analysis of phytochrome expression in transgenic lines. Seedlings were cultivated under the continuous white light (cW, 55 μmol photons m^-2^ s^-1^) for four days. The red number below each band represents the band intensity, which was quantified using ImageJ with normalization to RbcL. b, Phenotypes of HO1L and HO1S adults. Strains were cultivated under the cW condition on the MS medium for three weeks (upper) or under the short-day condition (SD, 8 h 55 μmol photons m^-2^ s^-1^ white light/16 h dark) on the Jiffy-7 soil for five weeks (lower). Scale bar, 5 mm.

From phenotypes of transgenic adults which were cultivated under the continuous white light for three weeks or under the short-day condition (SD, 8 h white light/16 h dark) for five weeks, it was clear to see that, leaves of both 7-1 *HO1L*/*gun2-1* and 4-5-3 *HO1S*/*gun2-1* were larger and greener than the *gun2-1* mutant (Fig. 7b). However, the 4-5-3 *HO1S*/*gun2-1* seedling was much larger than that of 7-1 *HO1L*/*gun2-1.* This result show that cytosolic HO1 can sustain total growth of Arabidopsis without supply of electron from the chloroplast ferredoxin.

### The HO1S transgenic line eliminates the *gun* phenotype

The expression of many *PhANGs* is dependent on the chloroplast functionality. Mutations in genes affecting the chloroplast function or treatments with NF or Lin can result in the strong down-regulation of many *PhANGs*. *gun* mutants including *gun2-1* that have a reduced ability to coordinate this nuclear response to chloroplast status were identified through the retention of *PhANGs* expression after an inhibitory NF treatment (36). The identification of a dominant *gun6-1D* mutant with increased FC1 activity led to the hypothesis that heme synthesized by FC1 is either the signal itself or a precursor of the signal (5).

To examine whether the *gun* phenotype is affected by *HO1L* or *HO1S* gene introduction, we examined the levels of representative *PhANGs*: *CARBONIC ANHYDRASE 1* (*CA1*), *LIGHT-HARVESTING CHLOROPHYLL-PROTEIN COMPLEX I SUBUNIT A4* (*LHCA4*), *LIGHT-HARVESTING CHLOROPHYLL A/B-BINDING PROTEIN 1.1* (*LHCB1.1*), *LHCB1.2*, *PHOTOSYSTEM II SUBUNIT QA* (*PSBQA*), and *RBCS1A*. With NF treatment, *PhANGs* expression levels in *gun2-1* are higher than those in Col-0 which is in accordance with previous reports (5, 36) (Fig. 8). However, the *gun* phenotype was not detected in both 7-1 *HO1L*/*gun2-1* and 4-5-3 *HO1S*/*gun2-1* transgenic lines (Fig. 8). It is of note that transcription levels of *PhANGs* in the 7-1 *HO1L/gun2-1* line were lower than those in the Col-0. From these data, it can be inferred that not only HO1L but also HO1S plays important roles in heme retrograde signal transduction.

**Figure 8.**
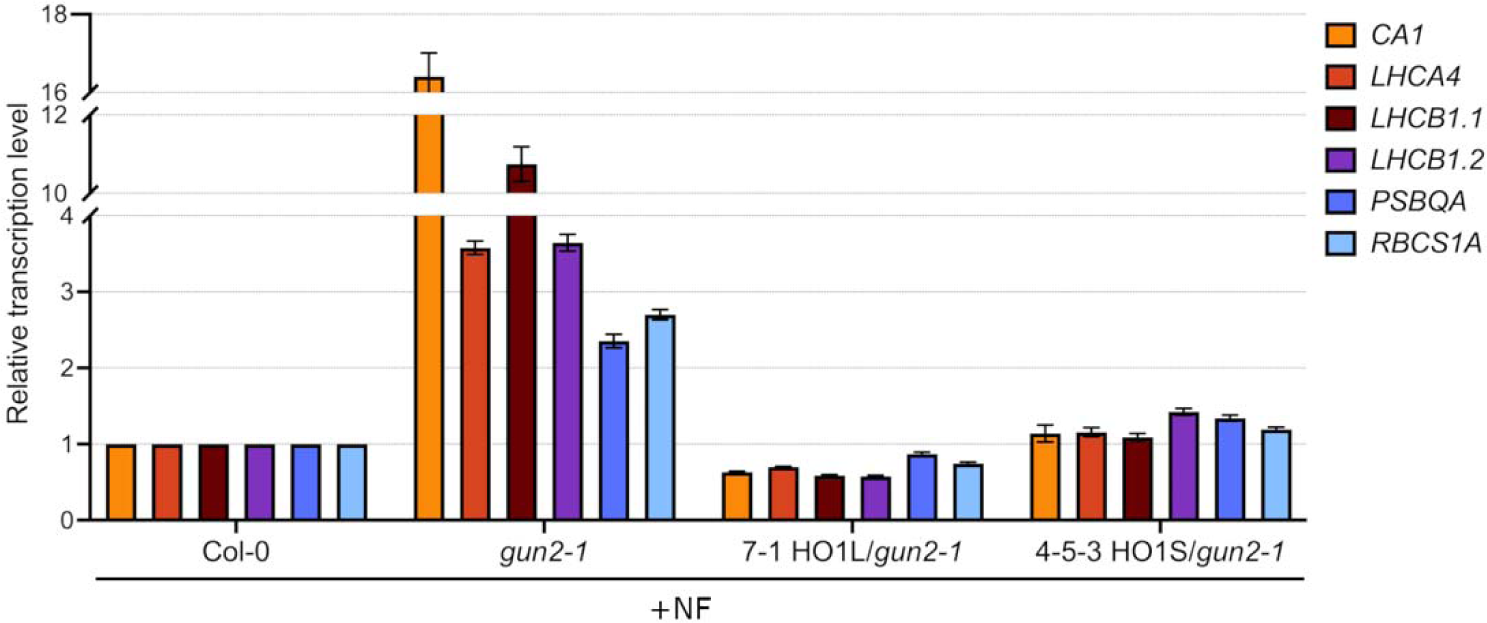
Relative transcription levels of *PhANGs* in NF-treated seedlings. Seedlings were grown under the continuous white light (cW, 55 μmol photons m^-2^ s^-1^) in MS medium supplemented with NF for four days. Primers used in qPCR are listed in *SI Appendix*, Table S1. At least three biological replicates were performed, and the representative one is shown.

## Discussion

### Alternative localization of HO1 in the plastid or cytosol by TSS regulation

This study showed that Arabidopsis *HO1* is regulated by TSS regulation in a light- and organ-dependent manner. Our study demonstrated that *HO1L* and *HO1S* products with or without the intact TP alternatively localize to plastids and the cytosol, respectively, and either form can function for heme catabolism and PΦB production in plant cells. Considering that the TSS regulation of *HO1* is observed in Arabidopsis and rice, the TSS-dependent alternative localization may be general in angiosperms. Since a single *HO* gene is present in *Marchantia polymorpha*, it is assumed that *HO* genes are diversified during the evolution of land plants. Currently, we do not know why the TSS regulation occurs specifically to *HO1* among HOs and the heme branch enzymes, but probably related to the predominant function of HO1 for PΦB biosynthesis. A comprehensive model is proposed to elucidate the regulatory pathway of *HO1* TSSs and the respective functions of their products (Fig. 9).

**Figure 9.**
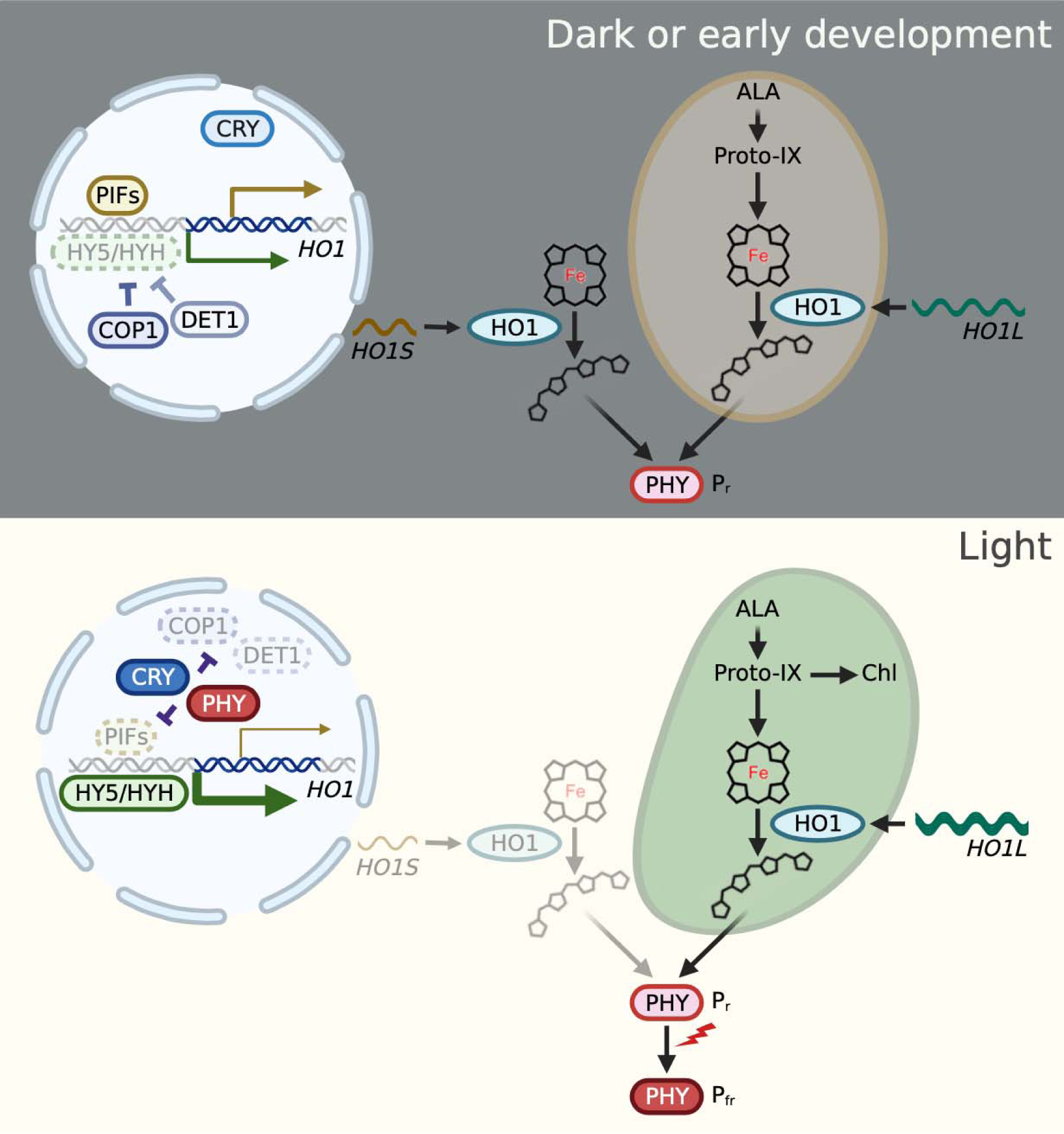
Proposed model of *HO1* TSS regulation and functions of *HO1L* and *HO1S* products. Under the dark or when seedlings are in the early development stage (upper), photomorphogenic transcription factors such as HY5 and HYH are targeted by COP1 complex with the help of DET1 complex for degradation. Skotomorphogenic transcription factors PIFs are presumed to control the transcription of *HO1*, downregulating the *HO1L* ratio and upregulating the *HO1S* ratio. Whereas under the light (lower), phytochrome is activated, transforms from the P_r_ form to the P_fr_ form, and migrates from the cytosol to the nucleus. Here it and the activated cryptochrome inhibit PIFs, COP1, and DET1, allowing HY5 and HYH to control the transcription of *HO1*, thereby *HO1L* ratio increases and *HO1S* ratio decreases. *HO1L* or *HO1S* product with or without the intact TP localizes to plastids or the cytosol, respectively. Significantly, not only *HO1L*, but also *HO1S* product functions in heme metabolism and PΦB generation. Light and dark blue boxes indicate inactivated and activated cryptochrome.

### Regulation of *HO1* TSSs by light signaling components

In the dark or during the early developmental stage of Arabidopsis, PIFs, known as skotomorphogenic transcription factors, repress the *PhANGs* expression (37). PIFs may also enhance the *HO1S* transcription. In contrast, photomorphogenic transcription factors HY5 and HYH are digested by 26S proteasome after ubiquitination with COP1 and DET1 complex (30). Therefore, the *HO1L* ratio remains low and the *HO1S* ratio remains significantly high in the dark or early developmental stage. Upon light absorption, phytochrome undergoes conformational changes and migrates from the cytosol to the nucleus (38), and cryptochrome is also activated. These photoreceptors mediate the inactivation of COP1, DET1, and PIFs, allowing HY5 and HYH to upregulate *HO1L* transcription, thereby increasing the *HO1L* ratio and decreasing the *HO1S* ratio. It is of note that *HO1* was included in the previous list of phytochrome-induced genes with alternative promoter selection (4), however, it was predicted that both *HO1L* and *HO1S* products were localized in the nucleus which is apparently incorrect.

According to the *in vivo* chromatin immunoprecipitation coupled with DNA chip hybridization (ChIP-chip) data, it has been determined that *HO1* is among the putative target genes of HY5 (39). However, ChIP-sequencing (ChIP-seq) and RNA-sequencing (RNA-seq) have identified that *HO1* is not included in the lists of direct target genes bound to PIF1/PIF3/PIF4/PIF5 (40). Further investigation is therefore necessary to confirm the direct interaction between HY5 and *HO1* promoter, and check whether PIFs directly bind *HO1* promoter or not. If not, it is required to determine precise mechanisms through which PIFs regulate the transcription of the *HO1* gene. Additionally, WRKY transcription factors represent one of the most extensive families in plants, comprising 72 members in Arabidopsis and over 100 in rice (41). WRKYs exhibit specific binding affinity to the TTGAC(C/T) W-box *cis*-element, located in the promoter region of their target genes (42). Moreover, the *HO1* promoter was characterized using *PlantCARE* (43), revealing the presence of three W-boxes located 1,798 bp upstream of the ATG starting codon. Therefore, it is crucial to explore whether WRKYs are involved in the regulation of *HO1* TSS.

### Involvement of HO1 in *gun*-dependent retrograde signaling

It is identified that the *HO1* mutant (*gun2-1*/*hy1-1*) showed a *gun* phenotype (36, 44). The discovery of a dominant *gun* mutant (*gun6-1D*) resulting in the overexpression of FC1 has led to a model in which FC1-derived heme mediates plastid signaling (5). Since the FC2-overexpressing line did not exhibit *gun* phenotypes, it is considered that increased flux of the FC1-derived heme may act as a positive signaling molecule that enhances the expression of *PhANGs*. It is hypothesized that in the *gun2-1* or *hy1-1* mutant, heme cannot be degraded, and resulting heme causes upregulation of *PhANGs* (45). Our complementation analysis of *gun2-1* showed either expression of HO1L or HO1S was efficient for complementing the *gun* phenotype (Fig. 8). This result clearly shows that heme degradation, either in plastid or cytosol, is effective for the recovery from the *gun* phenotype, supporting a heme signal hypothesis. Of note, since the expression of HO1L was more efficient for complementation of the *gun* phenotype, heme degradation in the plastid, the site of heme biosynthesis, is more effective than the cytosol where the distributed heme may be dispersed. To demonstrate this hypothesis, the micro-localization of heme should be determined since total and free heme levels are not correlated with the levels of *gun* phenotypes (46). Expression of fluorescent heme sensor protein (47) in each organelle and nucleus, which can monitor the micro-localization of heme, may solve this challenging question.

### Physiological significance of the cytosolic heme catabolic pathway

Heme catabolism is believed to occur entirely within the plastid, followed by the release of PΦB to the cytosol where holo-phytochrome assembly occurs (23, 48). It is reported that HO1 (22) and HY2 (23) are ferredoxin-dependent enzymes, requiring localization of both enzymes within the plastid. Meanwhile, by co-expressing apo-phytochrome and Arabidopsis *HO1* and *HY2* genes in *Pichia pastoris*, holo-phytochrome was successfully synthesized in the cytosol, suggesting that plastid ferredoxin is not a prerequisite as an electron donor for both enzymes (49). Actually, the Arabidopsis HO1 assay showed ∼20% activity in the absence of ferredoxin compared to the complete condition (22). Recently, a study showed that BV-IXα can be produced in the presence of high concentration of NADPH without ferredoxin (50). This study demonstrates that the cytosolic heme catabolic pathway predominantly functions during early and skotomorphogenic developments. We hypothesize that during such chloroplast undeveloped conditions, in which photosynthetic electron transport is not functioning, cytosolic reductants presumably NADPH may donate electrons to HO1 to proceed with PΦB biosynthesis. Once chloroplast is developed, the plastid pathway becomes significant as electrons driven by photosynthetic electron transport can be transferred to both enzymes in the chloroplast. Our complementation analysis of *gun2-1* (Fig. 5) showed more efficient growth recovery by *HO1S* than *HO1L* product, suggesting the critical role of the cytosolic heme catabolic pathway during early and skotomorphogenic developmental stages for plant growth. To our surprise, only cytosolic heme metabolism can sustain whole growth of Arabidopsis without electron supply from chloroplast ferredoxin.

One crucial question is whether cytosolic BV-IXα produced by HO1S returns to chloroplast for further production of PΦB by HY2. As we showed, HY2 is not regulated by promoter switching (*SI Appendix,* Fig. S1h). However, in Araport 1.1 of The Arabidopsis Information Resource (TAIR) (https://www.arabidopsis.org/index.jsp), we found that there are distinct splicing variants of *HY2* transcripts, of which the longer one containing the exon 1 having an intact chloroplast TP, while the shorter one lacking the exon 1 has no TP (*SI Appendix,* Fig. S5a). To check whether such short transcript of *HY2* is present or not, we designed the specific primers and analyzed by PCR (*SI Appendix,* Fig. S5b). We can detect the short *HY2* transcript in Col-0, *gun2-1*, and 4-5-3 *HO1S*/*gun2-1* transgenic line, suggesting that in addition to chloroplast, HY2 is also localized in the cytosol by alternative splicing. Currently, we do not know how this alternative splicing is regulated. In addition to the HY2 localization, this point will be clarified in the future.

Fig. 9 shows the model of TSS regulation of *HO1*. Currently, we are considering that the cytosolic heme catabolic pathway is important for holo-PHYA assembly during the early stage of development when plants undergo heterotrophic growth. In this stage, *HO1S* is highly transcribed by the TSS regulation, accumulating cytosolic HO1. We also presume that cytosolic HY2 is produced by alternative splicing, although the detailed regulation is not known. Upon illumination, *HO1S* transcription attenuates while that of *HO1L* is activated, resulting in the accumulation of HO1 within the plastid. At this stage, activated photosynthetic electron transport may give sufficient electrons via ferredoxin to HO1 and HY2, which facilitates consistent production of holo-PHYB necessary for light responses. Our results challenge the conventional sense of phytochrome biosynthesis and demonstrate the importance of a new phytochrome biosynthetic pathway in the cytosol.

## Materials and Methods

### Plant materials and growth conditions

Arabidopsis Col-0 and Col-0 based *phyA-211*, *phyB-9*, *phyA-211 phyB-9*, *hy5-215*, *hy5-2 hyh* (SALK_056405, Wisc DsLox 253D10), *pifq*, *cop1-4*, *det1-1*, and *gun2-1* were utilized in this research. Col-4, Col-4 based *cry1-304*, *cry2-1*, and *cry1-304 cry2-1*, Ler, and Ler based *hy1-1* were also utilized here. Arabidopsis seeds were sterilized and sown on the 2/3 Murashige and Skoog (MS) medium supplemented with 1% sucrose. Subsequently, they underwent a three-day cold treatment before being transferred to a temperature of 22 °C and exposed to white light (55 μmol photons m^-2^ s^-1^) for five hours, aimed at inducing germination, unless stated otherwise. For the treatment of NF or Lin, they were added directly to the MS medium at a final concentration of 5 μM or 220 μg/ml, respectively.

### Vector construction

Vectors used in this study were described in the Supplemental materials and methods for vector construction *(S1 Appendix*, Supporting text).

### TSS-seq, RNA extraction, qPCR, 5’ RACE-PCR, and 5’ RACE-qPCR

For TSS-seq screening, RNA was extracted from Arabidopsis and rice tissues, and subjected to TSS-seq using the Cap-Trapper method as described elsewhere (51).

Seedlings cultivated under various conditions were harvested and frozen immediately in the liquid nitrogen. Total RNA was extracted using the RNeasy Plant Mini Kit (QIAGEN) including the treatment with RNase-Free DNase Set (QIAGEN).

For normal qPCR analysis, 500 ng total RNA was reverse transcribed into cDNA with the PrimeScript RT Master Mix (TaKaRa). qPCR was performed using corresponding primers listed in *SI Appendix*, Table S1, with THUNDERBIRD Next SYBR qPCR Mix (TOYOBO) on a QuantStudio 1 Real-Time PCR System (ThermoFisher).

For 5’ RACE-PCR, 1 μg total RNA was reverse transcribed into cDNA with the SMARTer RACE 5’ Kit (Clontech). PCR was performed using primer GSP-*HO1* and the Universal Primer Mix (UPM, Clontech) with KOD One PCR Master Mix (TOYOBO) on a Light-Cycler 480 (BIO-RAD) (*SI Appendix,* Table S1). PCR products were separated on 2.5% agarose gels. Target bands were cut out, purified with the FavorPrep Gel/PCR Purification Kit (FAVORGEN), cloned into the pCR-Blunt II-TOPO vector (Invitrogen), and transformed into *Escherichia coli* DH5α. Plasmids were next extracted from transformed *E. coli* and target bands were confirmed by sequencing analysis. *HO1L* ratio (*HO1L*/*HO1*) and *HO1S* ratio (*HO1S*/*HO1*) were quantified with gel pictures using the CSAnalyzer4 software (ATTO).

To calculate *HO1L* and *HO1S* ratios more efficiently, qPCR was performed using two primer pairs, *HO1*-F and *HO1*-R, and *HO1L*-F and *HO1L*-R with THUNDERBIRD Next SYBR qPCR Mix on a QuantStudio 1 Real-Time PCR System (*SI Appendix,* Table S1). Plasmid pENTR/D-TOPO-*HO1L* linearized with restriction enzyme *Not*I (TOYOBO) was utilized for qPCR to draw standard curves of *HO1L* and total *HO1*. Copy numbers of *HO1L* and total *HO1* in the samples were calculated with measured Ct values from the standard curves. *HO1L* ratio was calculated by the formula: Ratio (*HO1L*)=*HO1L*/*HO1*=copy number (*HO1L*)/copy number (*HO1*). *HO1S* ratio was calculated by the formula: Ratio (*HO1S*)=*HO1S*/*HO1*=1–Ratio (*HO1L*).

### Transient expression assay

Vectors encoding *p35S*::*HO1L*-*mClover3* and *p35S*::*HO1S*-*mClover3*, along with *AtRBCS1A-TP*-*mScarlet*, or *mScarlet* alone, were transiently transformed into leaf cells of pea and tobacco using biolistic bombardment, as previously described (52), in order to determine subcellular localization using confocal laser-scanning microscopy (CLSM).

### Agrobacterium-mediated floral dip transformation

Vectors encoding *p35S*::*HO1L*-*mClover3* and *p35S*::*HO1S*-*mClover3* were transformed into *Agrobacterium tumefaciens* GV3101, respectively according to the freeze-thaw method (53). After that, GV3101 strain containing vectors were transformed into *gun2-1* or *hy1-1* with the Agrobacterium-mediated floral dip transformation (54). T3 or T4 generations were used for this study.

### Fluorescence microscopic analysis

As for subcellular localization analysis with leaf cells of pea and tobacco, HO1L-mClover3 and HO1S-mClover3 were viewed with a TCS SP5X CLSM (Leica Microsystems). The mClover3 fusions and Chl were excited with an argon laser line (488 nm) to detect fluorescence at 500-530 nm and 680-700 nm, respectively, while mScarlet and mScarlet fusion were excited with a He/Ne laser line (543 nm) to view fluorescence at 565-615 nm.

As for protoplasts released from well-expanded leaves, HO1L-mClover3 and HO1S-mClover3 were detected under the CLSM (FV3000, OLYMPUS). Excitation wavelength was 488 nm, and emission wavelength was 500-550 nm for the mClover3 and 662-691 nm for the Chl, respectively.

### Immunoblot analysis

Four-day-old seedlings were harvested and frozen immediately in the liquid nitrogen. After being crushed, debris was denatured in the Laemmli buffer. Protein amounts were measured by RC DC Protein Assay (BIO-RAD). In order to detect HO1, 10 μg protein was separated using 10% SDS-PAGE and subsequently immunoblotted with an anti-HO1 antibody. For detecting PHYA and PHYB, 10 μg protein was separated in 8% SDS-PAGE and immunoblotted with anti-PHYA and anti-PHYB antibodies.

### Phenotypic analysis

Photographs were taken of four-day-old seedlings after they were transferred to the 0.7% agarose plates. Hypocotyl lengths were determined by analyzing captured photographs using the NIH ImageJ software.

Four-day-old strains were harvested, weighted, and immersed in 1 ml 80% acetone overnight to extract Chl and carotenoid (55). Debris was removed by centrifugation at 18,000 g for 5 min. The absorption was read with a spectrophotometer (JASCO V-730) at 663, 647, and 470 nm. Chl *a* amount was calculated using the formula: Chl *a* (mg g FW^-1^) =(12.25×A663−2.79×A647)/1000/m (fresh weight of seedlings). Chl *b* amount was calculated using the formula: Chl *b* (mg g FW^-1^)= (21.50×A647−5.10×A663)/1000/m. Carotenoid amount was calculated using the formula: carotenoid (mg g FW^-1^)=(1000×A470−1.80×Chl *a*−85.02×Chl *b*)/198/1000/m. Heme was extracted and measured from the samples according to the published protocol (46).

## Acknowledgments

We sincerely thank Prof. Nobuyoshi Mochizuki (Graduate School of Science, Kyoto University) for providing mutant lines including *phyA-211*, *phyB-9*, *phyA-211 phyB-9*, *cop1-4*, and *det1-1*, Prof. Koichi Kobayashi (Graduate School of Science, Osaka Metropolitan University) for providing the *hy5-215* mutant line, Prof. Takayuki Kohchi (Graduate School of Biostudies, Kyoto University) for providing the anti-HO1 antibody, Prof. Tsuyoshi Nakagawa (Interdisciplinary Center for Science Research, Shimane University) for providing Gateway destination vectors including pUGW0, pUGW2, pGWB401, pGWB601, and pGWB602, Dr. Noriyuki Suetsugu (Graduate School of Arts and Sciences, The University of Tokyo) and Prof. Matthew J. Terry (School of Biological Sciences, University of Southampton and Institute for Life Sciences, University of Southampton) for suggestions and comments concerning the research. This work was supported by The Sasakawa Scientific Research Grant.

## Notes

### Competing Interest Statement

The authors have declared no competing interest.

