## Supplemental files for "Cytosolic heme metabolism by alternative localization of heme oxygenase 1 in plant cells"

Tatsuru Masuda

**This PDF file includes:**

Supporting text

Figures S1 to S5

Table S1

SI References

Supporting Information Text

Materials and Methods

Vector construction

*HO1* transcripts, specifically *HO1L* and *HO1S*, obtained from Col-0 cDNA, were cloned into the pENTR/D-TOPO vector (Invitrogen) following the manufacturer's instructions, resulting in the formation of pENTR/D-TOPO-*HO1L* and pENTR/D-TOPO-*HO1S*. Gateway destination vectors tagged with *mClover3* (1) or *mScarlet* (2) were constructed as follows.

An *mClover3* fragment was amplified by PCR using primers, GW-*3xGGGGS*-F, *3xGGGGS*-F, *3xGGGGS-mClover3*-*C*-F, *mClover3*-R, and *XFP-C*-GW-R, from plasmid pKanCMV-*mClover3*-*mRuby3* (deposited by Michael Lin; Addgene #74252) (1). It was inserted into the *Afe*I site of pUGW2 (3) to generate an intermediate vector pGWCL3 by the SLiCE method (4). An *Xba*I-Gateway cassette-*3xGGGGS*-*mClover3*-*Sac*I fragment from pGWCL3 amplified by PCR using a pair of primers, *Xba*I-GW-F and GW-*Sac*I-R, was inserted into *Xba*I and *Sac*I sites of pGWB602 vector (5) by the Hot Fusion method (6) to create a Gateway-compatible destination vector pB6GWCL.

As for *mScarlet* destination vectors, a *Hind*III-*p35S*-Gateway cassette-*Sac*I fragment from pUGW0 (3) was ligated into *Hind*III and *Sac*I sites of pGWB402 (3) to generate an intermediate vector pGWB402N. A DNA fragment of the promoter sequence of *AtUBQ10* gene (At4g05320) amplified by PCR using primers, *pUbi10-B4*-F, *pUbi10-B4*-R, *attB4*-*adapt*-F, and *attB1R*-*adapt*-R was cloned into pDONR P4-P1R (Invitrogen) to construct a plasmid pDONR P4-P1R/*pAtUbi10*. To construct a Gateway destination vector under the control of *pAtUBQ10*, DNA fragments of *pAtUBQ10* for N- and C-terminal tags (*pUBQ10N* and *pUBQ10C*) were obtained from the pDONR P4-P1R/*pAtUbi10* by PCR using primers *pUBQ10*-F, *pUBQ10N*-R, and *pUBQ10C*-R. *pUBQ10N* was inserted into *Hind*III and *Xba*I sites of pGWB402N by the Hot Fusion to create a pB4UxGW vector, while *pUBQ10C* was inserted into the same sites of pGWB402 and pGWB601 (5) to make pB4UGWx and pB6UGWx vectors, respectively. Fragments of *mScarlet*-*3xGGGGS* and *3xGGGGS*-*mScarlet* amplified by PCR from a plasmid pRD134 (deposited by Michael Lin; Addgene #74252) (2) using primers, GW-*mScarlet-N*-F, *XFP-3xGGGGS*-R, *3xGGGGS*-R, and *3xGGGGS*-GW-R for *mScarlet-3xGGGGS*, and GW-*3xGGGGS*-F, *3xGGGGS*-F, *3xGGGGS-mScarlet-C*-F, and *XFP-C*-GW2-R for *3xGGGGS*-*mScarlet*, were inserted into *Afe*I site of pB4UxGW and pB4UGWx to create Gateway-compatible destination vectors, pB4UScGW (for N-terminal *mScarlet* fusion) and pB4UGWSc (for C-terminal *mScarlet* fusion), respectively. An *Sac*I-Gateway cassette-*3xGGGGS*-*mScarlet*-*Sac*I fragment from pB4UGWSc by PCR using a pair of primers, *Sac*I-*pUBQ10*-F and GW-*Sac*I-R, was inserted into *Sac*I sites of the pB6UGWx by the Hot Fusion to create a Gateway-compatible destination vector, pB6UGWSc (for C-terminal fusion).

Gateway entry clones such as pENTR/D-TOPO-*HO1L* and pENTR/D-TOPO-*HO1S* were subjected to an LR reaction (Invitrogen) with pB6GWCL to construct the corresponding expression vectors, pB6GWCL/*p35S*::*HO1L-mClover3*:*nopaline synthase terminator* (*tNOS*) and pB6GWCL/*p35S*::*HO1S-mClover3*:*tNOS*. As for the chloroplast marker, plasmid harboring *AtRBCS1A-TP* for chloroplastidic stromal marker (Pt/pDONR221) (7) was subjected to an LR reaction with pB6UGWSc to construct the corresponding expression vector pB6UGWSc/*pAtUBQ10*::*AtRBCS1A-TP*-*3xGGGGS-mScarlet*:*tNOS*. Nucleotide sequences of the inserted genes in all the plasmids described above were confirmed by sequencing analysis. PCR primers were listed in *SI Appendix*, Table S1.

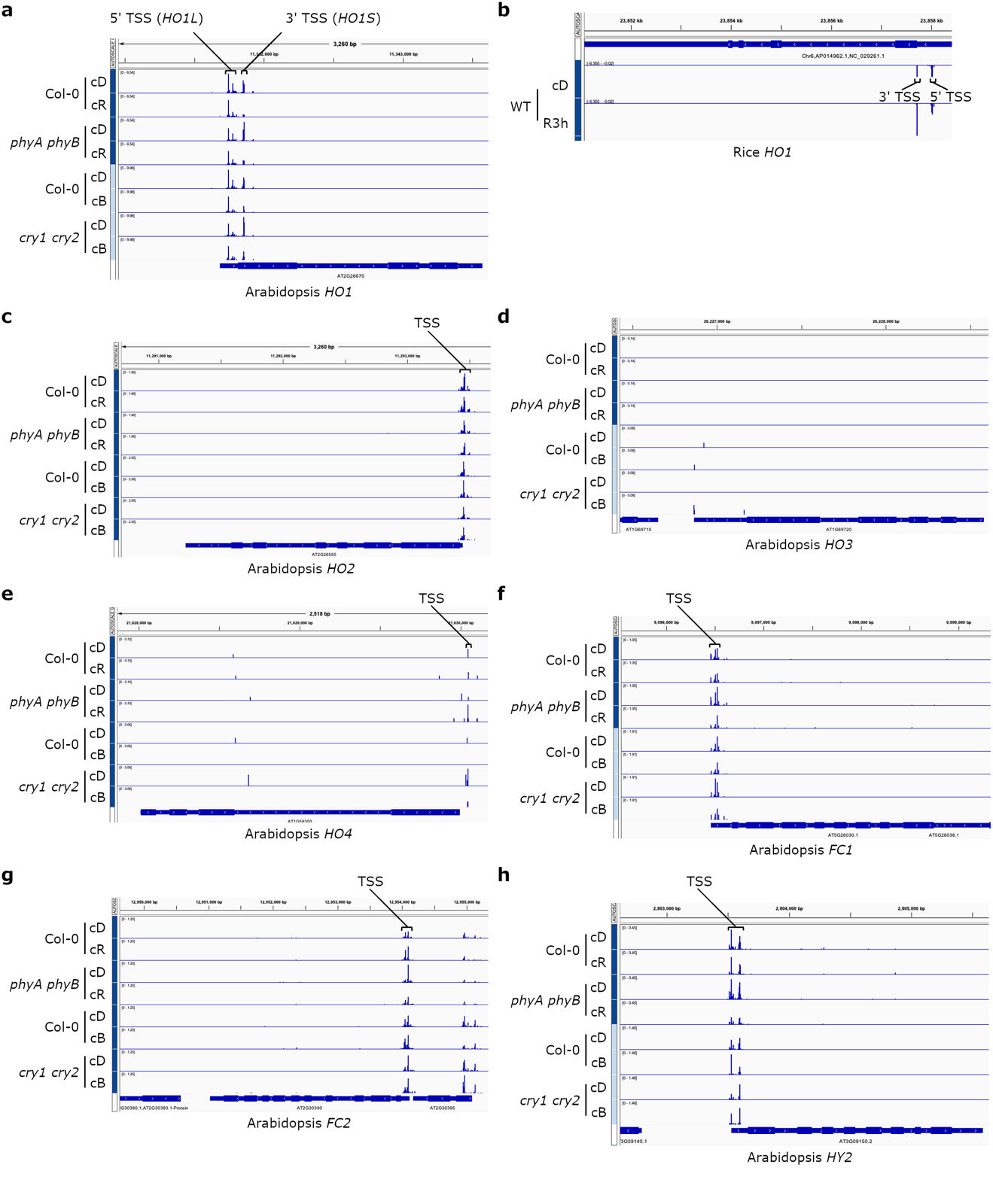

Fig. S1. CAGE-seq analysis of TSS of genes functioning in heme synthesis and metabolism branch. Screenshots of TSS signals of genes include *HO1* (a), *HO2* (c), *HO3* (d), *HO4* (e), *FC1* (f), *FC2* (g), and *HY2* (h) of Arabidopsis, and *HO1* (b) of rice. Arabidopsis Col-0, and Col-0 based *phyA phyB* and *cry1 cry2* mutants were cultivated under the continuous dark (cD), red light (cR) or blue light (cB) for four days. The wide type (WT) of rice was cultivated under the cD for four days or first under the cD for four days and then illuminated under the red light for three hours (R3h).

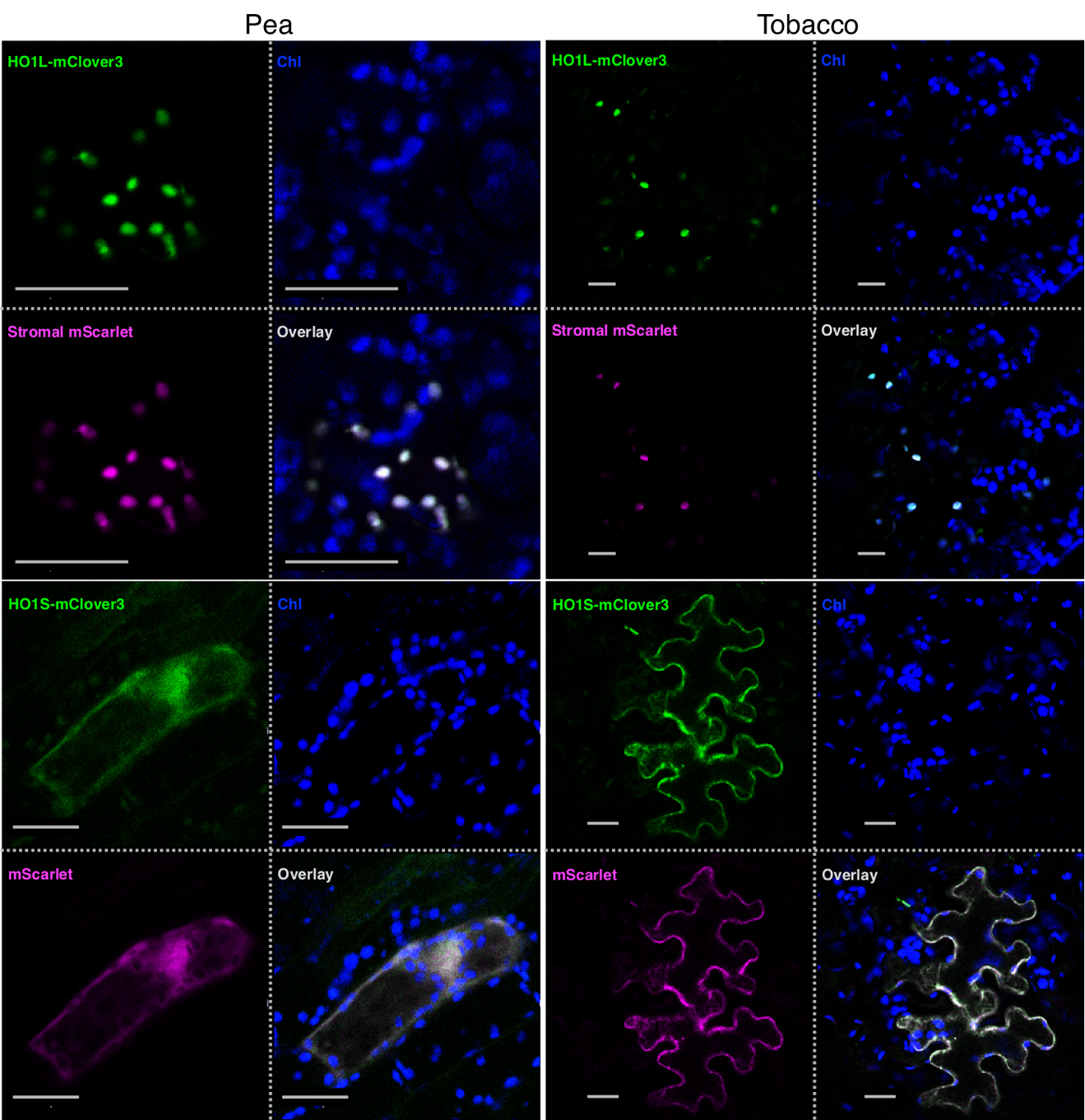

Fig. S2. Transient expressions of HO1L-mClover3 or HO1S-mClover3 in plant leaves. Pea (left) and tobacco (right) were utilized in the transformation. HO1L-mClover3 and HO1S-mClover3 were viewed with CLSM. *At*RBCS1A-TP-mScarlet served as a chloroplast-stromal marker. mScarlet alone was generally dispersed in the cytosol. Single-section images of the GFP channel (upper left), Chl channel (upper right), RFP channel (lower left), or their overlay (lower right) in each panel are shown. Bars indicate 20 μm.

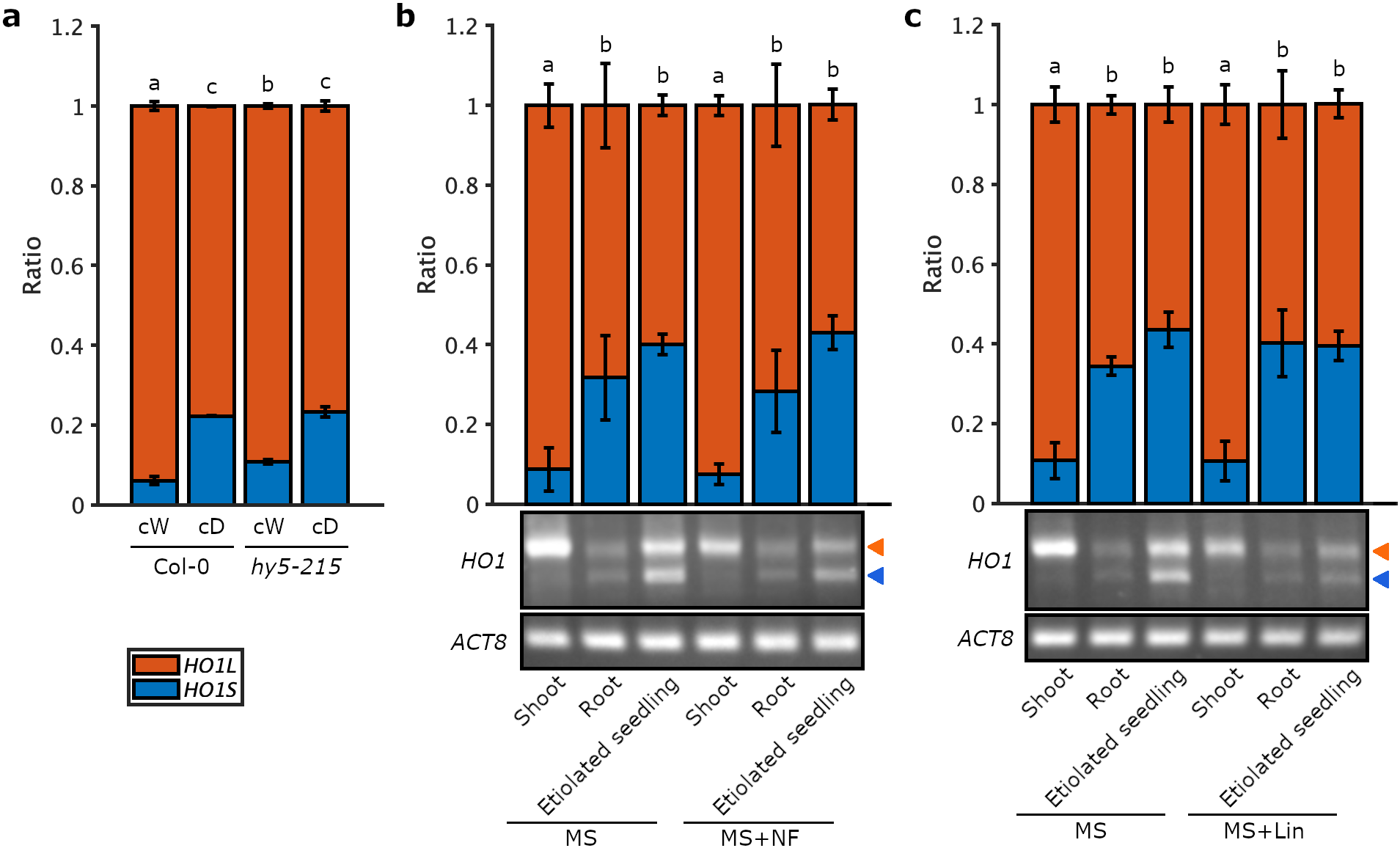

Fig. S3. 5’ RACE-qPCR and 5’ RACE-PCR analysis of *HO1* TSSs in transcription factor mutant and herbicide treated seedlings. a, 5’ RACE-qPCR analysis of *HO1L* and *HO1S* ratios in the *HY5* mutant. Col-0 and *hy5-215* were cultivated under the continuous white light (cW, 55 μmol photons m^-2^ s^-1^) or dark (cD) for four days. b and c, 5’ RACE-PCR analysis of *HO1L* and *HO1S* ratios in NF or Lin treated Col-0. Col-0 was cultivated under the cW or cD condition for five days. 5 μM NF (b) or 220 μg/ml Lin (c) was added to the MS medium.

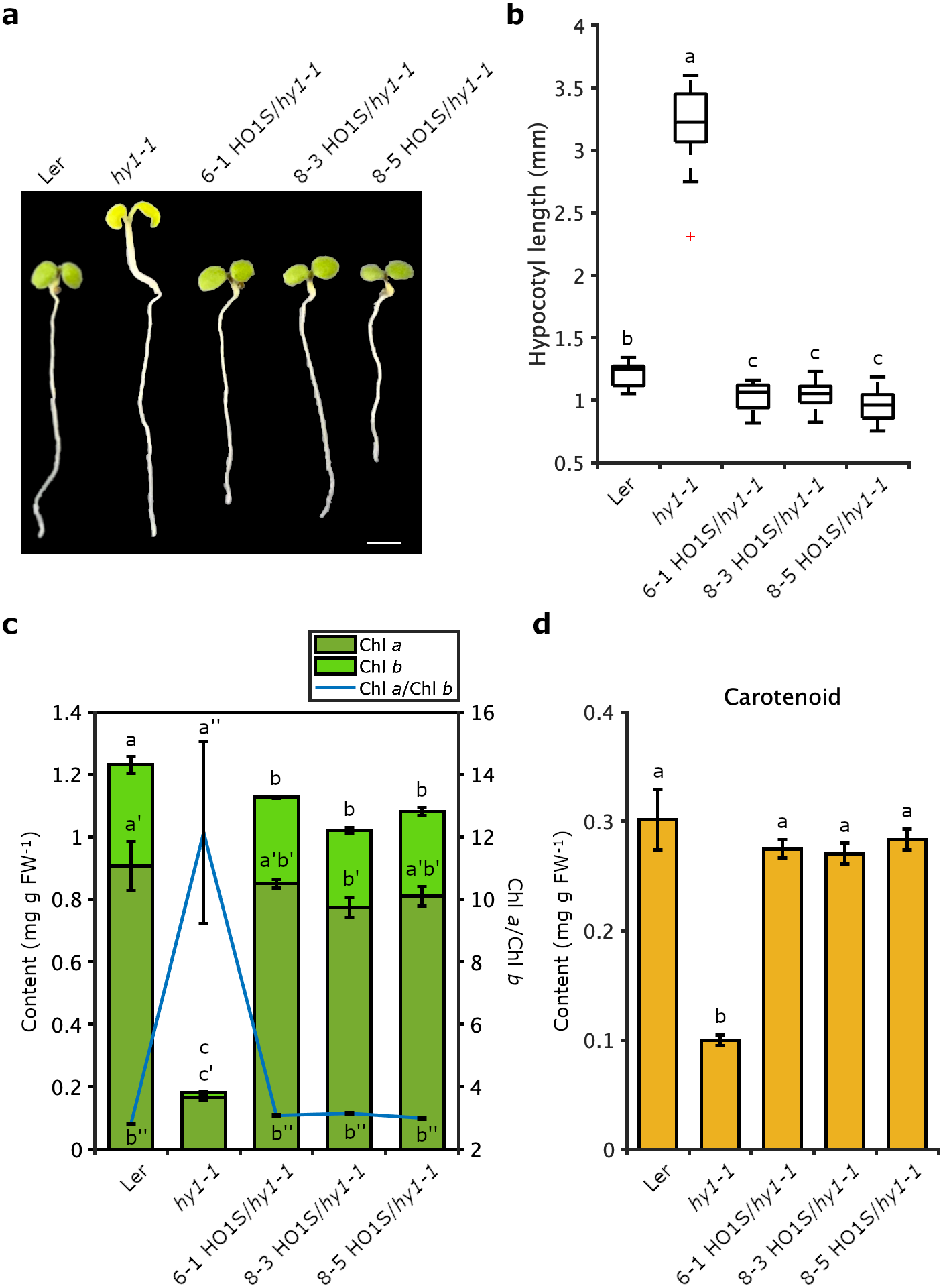

Fig. S4. Hypocotyl length and pigment contents of HO1S/*hy1-1* transgenic lines. a, Phenotypes of four-day-old HO1S/*hy1-1* transgenic lines. Seedlings were cultivated under the continuous white light (cW, 55 μmol photons m^-2^ s^-1^) for four days. Scale bar, 2 mm. b, Hypocotyl length of HO1S/*hy1-1* transgenic lines. Strains were cultivated under the cW condition for four days. Hypocotyl lengths were measured using ImageJ, and measurements were taken for 20 seedlings from each line. p<0.05, one-way ANOVA, and Turkey’s multiple comparison tests. c and d, Chl content (c), Chl *a*/Chl *b* ratio (c), and carotenoid content (d) of HO1S/*hy1-1* transgenic lines. Seedlings were grown under the cW condition for four days. The data mean ± SD was calculated from values of three technical replicates.

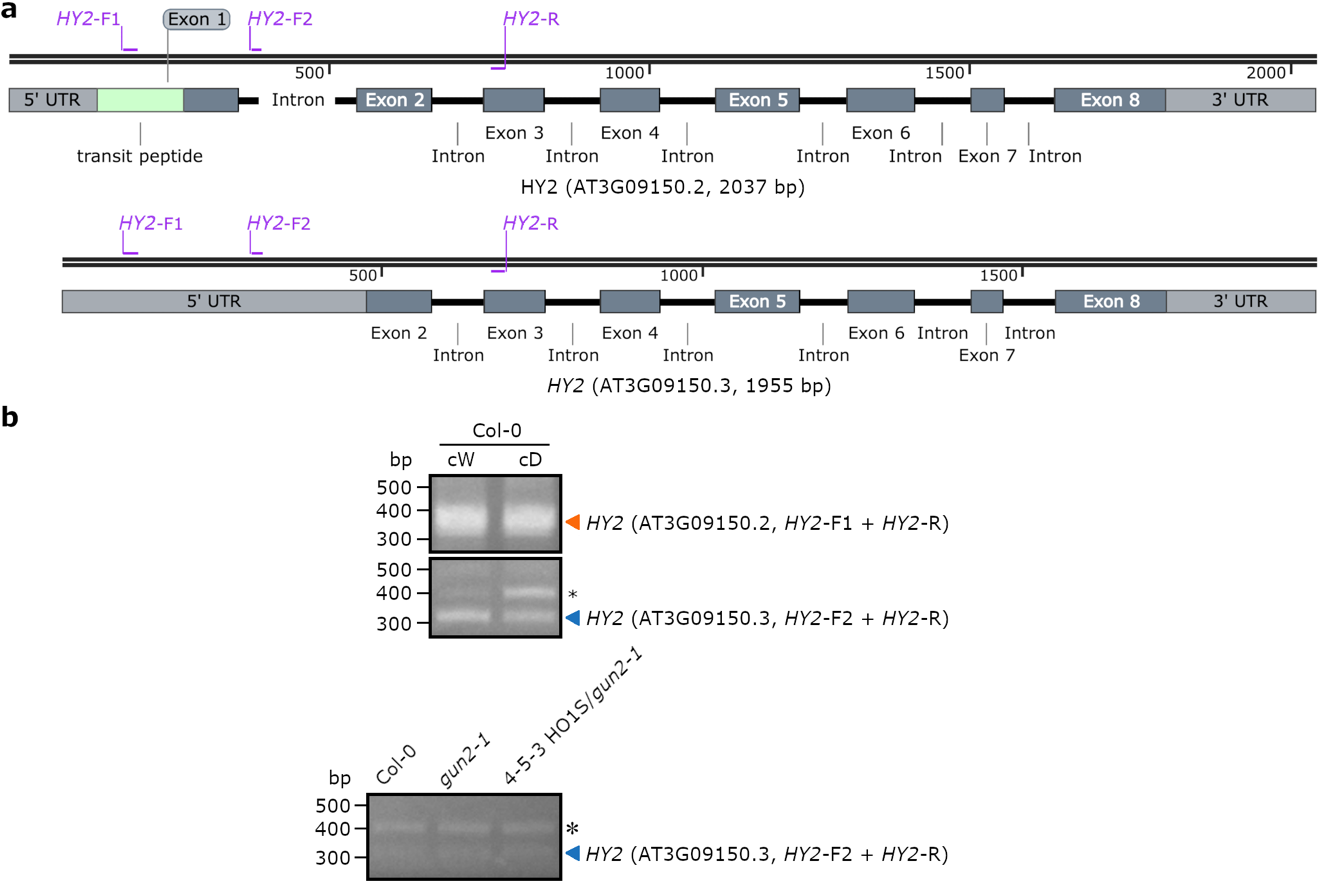

Fig. S5. The identification of *HY2* transcripts. (a) *HY2* pre-mRNA structures, AT3G09150.2 and AT3G09150.3. Light grey box illustrates 5’ UTR and 3’ UTR. Dark grey box illustrates exon with light green box illustrating TP. Black line illustrates intron. Primers are illustrated above gene structures. The diagram was constructed with SnapGene software based on the data from The Arabidopsis Information Resource (TAIR) (<https://www.arabidopsis.org/>) and UniProt. (b) DNA gel electrophoresis of *HY2* PCR products. Primer pair *HY2*-F1 and *HY2*-R was utilized for the detection of AT3G09150.2 transcript, and primer pair *HY2*-F2 and *HY2*-R was utilized for the detection of AT3G09150.3 transcript. Four-day-old seedlings grown under continuous white light (cW, 55 μmol m^-2^ s^-1^) or dark (cD) were harvested for mRNA extraction. 5’ RACE was utilized for the reverse transcription (RT) of upper panel and normal RT was performed in the lower panel. Orange and blue arrowheads illustrate target fragments of AT3G09150.2 and AT3G09150.3, respectively. Star illustrates the genome DNA. Fragments were checked by cloning and sequencing.

Table S1. Primers.

| Name | Sequence (5’-3’) |
| --- | --- |
| *3xGGGGS*-F | GCTGGTGGAGGCGGTTCAGGCGGAGGTGGCTCTGGCGGTGGCGGATCG |
| *3xGGGGS-mClover3-C-*F | GGCGGTGGCGGATCGATGGTGAGCAAGGGCGAGGA |
| *3xGGGGS-mScarlet-C*-F | GGCGGTGGCGGATCGATGGTGAGCAAGGGCGAGGCA |
| *3xGGGGS-*GW-R | TACAAACTTGTGATAACAGCGCTTCCGCCACCGCC |
| *3xGGGGS-*R | GCCTGAACCGCCTCCACCCTTGTACAGCTCGTCCATGCCG |
| *35S*-F | TCCACTGACGTAAGGGATGACGCACAATC |
| *ACT8*-F | ACTGTGCCTATCTACGAGGGTTTC |
| *ACT8*-R | CCCGTTCTGCTGTTGTGGT |
| *attB1R*-adapt-R | GGGGACTGCTTTTTTGTACAAACTTG |
| *attB4*-adapt-F | GGGGACAACTTTGTATAGAAAAGTTG |
| *CA1*-F | GAGAAATACGAAACCAACCCT |
| *CA1*-R | ACATAAGCCCTTTGATCCCA |
| GSP-*HO1* | CGAGTATCCGCTCTGCCACCTTTCTGCC |
| GW-*3xGGGGS*-F | TGGTTGATAACAGCGCTGGTGGAGGCGGT |
| GW-*mScarlet-N*-F | ATTAACAGTCTAGAATGAGCATGGTGAGCAAGGGCGAGGCA |
| GW-*Sac*I-R | CGATCGGGGAAATTCGAGCTCTAAGCCTTGTACAGCTCGT |
| *HO1*-F | AGTCGCCGTCTTTAGTGGTG |
| *HO1*-R | ACCTTCAACAGTAGGTTCCCA |
| *HO1L*-F | GCGTATTTAGCTCCGATTTCTTCA |
| *HO1L*-R | TCCTAGGTCGAAGGAAAAGTG |
| *HY2-*F1 | TGCTTCAAGGCACCAAACCCACCTGT |
| *HY2-*F2 | TCGTTAGTGTAGTGGGAGGA |
| *HY2-*R | ACTCAGGCTCCATGAAACCCGCAAA |
| *LHCA4*-F | AACCCGCTTAACTTTGCTCCTAC |
| *LHCA4*-R | CAAACCCTAAGAATGCCAACATC |
| *LHCB1.1*-F | GAGCCAAGTTCTATCTGTTTG |
| *LHCB1.1*-R | TCTACCATCCACCACAAACAC |
| *LHCB1.2*-F | GATGGGAGCTGTTGAAGGCT |
| *LHCB1.2*-R | CCTCTGGGTCGGTAGCAAGA |
| *mClover3-R* | CTTGTACAGCTCGTCCATGCCATGTGTAATCCCGGCGGCGGTCACGAA |
| *PSBQA*-F | AATGGCTCTGGAAGAGTGGC |
| *PSBQA*-R | AATAGCATCGGCGAGGACAG |
| *pUbi10-B4*-F | ATAGAAAAGTTGTTTCAGTAATAAACGGCGTCAAAG |
| *pUbi10-B4*-R | TTTGTACAAACTTGCCATCTGTTAATCAGAAAAACTCAG |
| *pUBQ10*-F | CGACGGCCAGTGCCAAGCTTTCAGTAATAAACGGCGTCAA |
| *pUBQ10C*-R | AACTTGTTGATAACTCTAGACTGTTAATCAGAAAAACTCAGATTAATCGA |
| *pUBQ10N*-R | ATAACAGCGCTCATTCTAGACTGTTAATCAGAAAAACTCAGATTAATCGA |
| *RBCS1A*-F | TTCCTGACCTTACCGATTCCG |
| *RBCS1A*-R | CGGTACACAAATCCGTGCT |
| *Sac*I-*pUBQ10*-F | CTATAAAACAATACCCAAAGAGCTCTTCTTCT |
| *Xba*I*-*GW-F | AGAACACGGGGGACTCTAGAGTTATCAACAAGTTTGTACAAAAAAGCT |
| *XFP-3xGGGGS*-R | GCCTGAACCGCCTCCACCCTTGTACAGCTCGTCCATGCCG |
| *XFP-C-*GW-R | ATTCGAGCTCTAAGCCTTGTACAGCTCGTCCATGC |
| *XFP-C-*GW2-R | GGGAAATTCGAGCTCTAAGCCTTGTACAGCTCGTCCATGCCG |

SI References

| 1. | B. T. Bajar *et al.*, Improving brightness and photostability of green and red fluorescent proteins for live cell imaging and FRET reporting. *Sci. Rep.* 6, 1–12 (2016). |
| --- | --- |
| 2. | D. S. Bindels *et al.*, mScarlet: a bright monomeric red fluorescent protein for cellular imaging. *Nat. Methods* 14, 53–56 (2017). |
| 3. | T. Nakagawa *et al.*, Development of series of gateway binary vectors, pGWBs, for realizing efficient construction of fusion genes for plant transformation. *J. Biosci. Bioeng.* 104, 34–41 (2007). |
| 4. | K. Motohashi, A simple and efficient seamless DNA cloning method using SLiCE from Escherichia coli laboratory strains and its application to SLiP site-directed mutagenesis. *BMC Biotechnol.* 15, 47 (2015). |
| 5. | S. Nakamura *et al.*, Gateway binary vectors with the bialaphos resistance gene,*bar*, as a selection marker for plant transformation. *Biosci. Biotechnol. Biochem.* 74, 1315–1319 (2010). |
| 6. | C. Fu, W. P. Donovan, O. Shikapwashya-Hasser, X. Ye, and R. H. Cole, Hot fusion: An efficient method to clone multiple DNA fragments as well as inverted repeats without ligase. *PLoS One* 9, e115318 (2014). |
| 7. | M. Aboulela *et al.*, Development of an R4 dual-site (R4DS) gateway cloning system enabling the efficient simultaneous cloning of two desired sets of promoters and open reading frames in a binary vector for plant research. *PLoS One* 12, e0177889 (2017). |
